# Postglacial recolonization of the Southern Ocean by elephant seals occurred from multiple glacial refugia

**DOI:** 10.1101/2024.11.18.622576

**Authors:** Andrew A. Berg, Megan Askew, Frederik V. Seersholm, Alexander J. F. Verry, A. Rus Hoelzel, Andreanna Welch, Karen Greig, Richard Walter, Michael Knapp, Axel Barlow, Johanna L.A. Paijmans, Jonathan M. Waters, Michael Bunce, Kate McDonald, Sue O’Connor, Brenda Hall, Paul Koch, Carlo Baroni, Maria Cristina Salvatore, Patrick Faulkner, Simon Y. W. Ho, Nicolas J. Rawlence, Mark de Bruyn

**Author notes:** **Corresponding Authors:** Mark de Bruyn, Australian Research Centre for Human Evolution, School of Environment and Science, Griffith University, Nathan QLD 4111, Australia., Nicolas J. Rawlence, Otago Palaeogenetics Laboratory, Department of Zoology, University of Otago, Dunedin, New Zealand. These authors contributed equally and should be considered joint first authors (Andrew A. Berg, Megan Askew). These authors contributed equally and should be considered joint last authors (Nicolas J. Rawlence, Mark de Bruyn). **Competing Interest Statement:** No competing interests.

## Abstract

The Southern Ocean is warming more rapidly than other parts of our planet. How this region’s endemic biodiversity will respond to such changes can be illuminated by studying past events, through genetic analyses of time-series data sets including historic and fossil remains. Archaeological and subfossil remains show that the southern elephant seal (*Mirounga leonina*) was common along the coasts of Australia and New Zealand in the recent past. This species is now mostly confined to sub-Antarctic islands and the southern tip of South America. We analysed ancient seal samples from Australia (Tasmania), New Zealand, and the Antarctic mainland to examine how southern elephant seals have responded to a changing climate and anthropogenic pressures during the Holocene. Our analyses show that these seals formed part of a broader Australasian lineage, comprising seals from all sampled locations from the south Pacific sector of the Southern Ocean. Our study demonstrates that southern elephant seal populations have dynamically altered both range and population sizes under climatic and human pressures, over surprisingly short evolutionary timeframes for such a large, long-lived mammal.

**Significance Statement:** Genetic data, alongside historic, archaeological, and subfossil remains show that Australasian populations of the southern elephant seal have been shaped by range expansions and contractions following the Last Glacial Maximum, with subsequent contractions during the late Holocene. These expansion and contraction events are likely to have been a direct result of climate change-induced habitat expansion and contraction, along with Indigenous and European sealing. Prehistoric climate change and more recent human pressures have substantially altered the geographic distribution and population size of southern elephant seals over short evolutionary timescales.

## Introduction

Polar and sub-polar species are particularly vulnerable to the impacts of global climate change due to their specialized habitats and the rapid warming experienced at high latitudes. The Arctic and Antarctic regions are warming at rates significantly higher than the global average, leading to drastic reductions in sea ice cover, altered food web dynamics, and shifts in species distributions (Hoegh-Guldberg & Bruno, 2010; Cole et al., 2019). Species such as polar bears (*Ursus maritimus*), Adélie penguins (*Pygoscelis adeliae*), and Antarctic krill (*Euphausia superba*) have shown variable responses to these environmental changes, including range shifts, changes in population size, and altered breeding patterns (Vianna et al., 2020; Fraser et al., 2012). Characterizing the resilience and adaptability of these species is critical for understanding broader ecosystem responses to warming climates. Studying how polar and sub-polar fauna have historically responded to past climate change, through genetic and archaeological evidence, can provide valuable insights into their potential future responses and guide conservation efforts in rapidly changing environments (Fraser et al., 2009; Gonzalez-Wevar et al., 2018).

Top predators play a critical role in the Antarctic ecosystem, yet are highly vulnerable to the impacts of climate change on any or all parts of the food chain, and the physical environment (e.g. Cole et al., 2019; Vianna et al., 2020). Southern elephant seals (*Mirounga leonina*) are a key species to test for the impacts of climate change and human impact due to the geographic and temporal range of individuals in the Late Quaternary. The Southern elephant seal plays a key role in the Antarctic ecosystem. The largest pinniped, and the largest non-cetacean marine mammal, southern elephant seals are apex predators that consume massive amounts of biomass to maintain their ∼4,000 kg bulk (males) during a nine-month foraging period (Gales et al., 1989; McMahon et al., 2005). This species inhabits a region that is undergoing rapid climate change – the sub-Antarctic and the Southern Ocean (Hoegh-Guldberg & Bruno, 2010). The present-day population is split into four major breeding ‘stocks’ distributed in and around the Southern Ocean: the South Georgia and Península Valdés stocks, both located in the Atlantic Ocean sector; the Kerguelen and Heard Islands stock, in the Indian Ocean sector; and the Macquarie Island stock in the Pacific Ocean sector (Figure 1). About 98% of the global population of southern elephant seals breed in these locations (McMahon et al., 2005).

**Figure 1.**
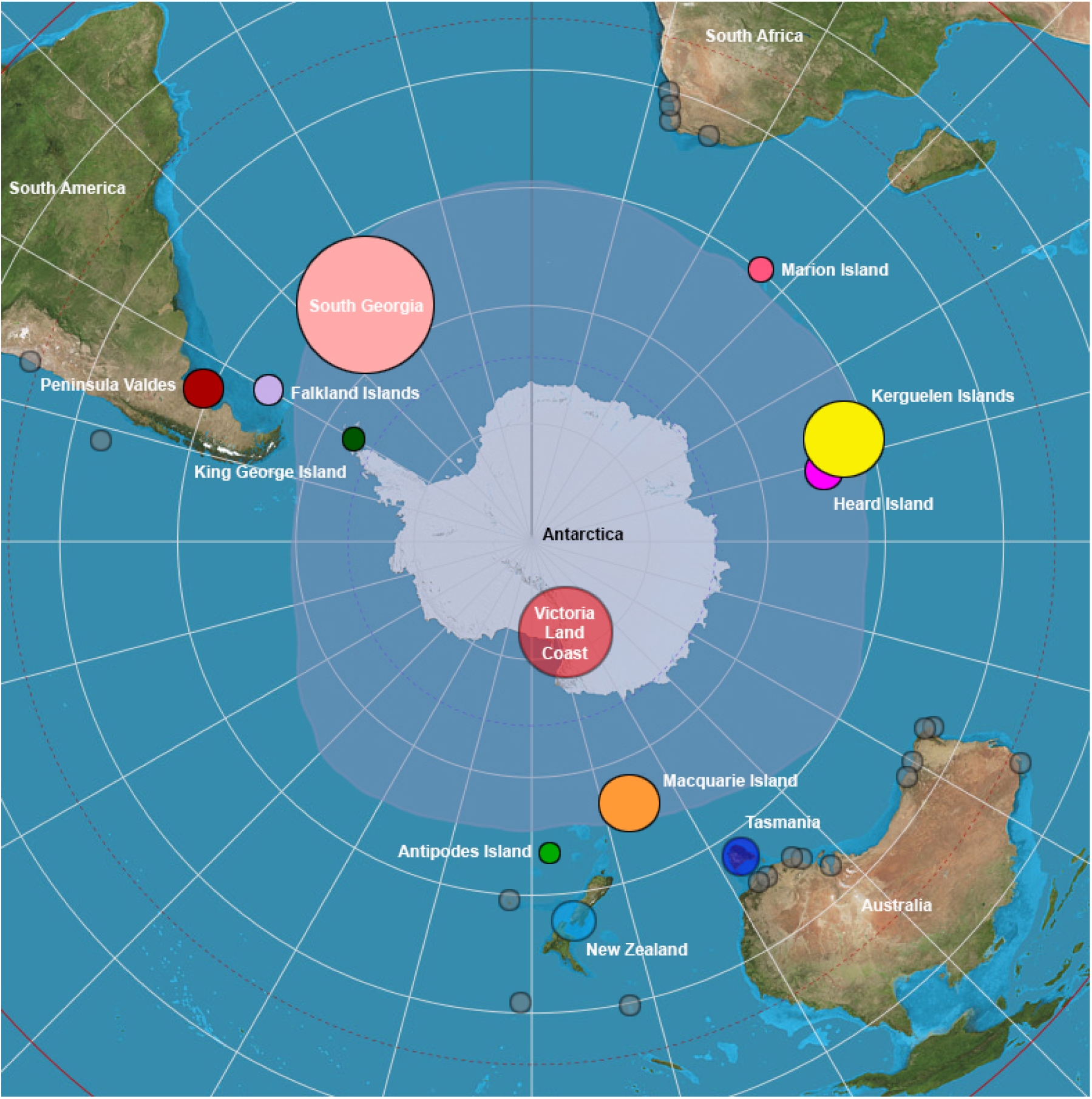
Map of known extant and extinct southern elephant seal distribution. Marker size is representative of the relative number of seals at each location. The map shows extant breeding colonies (solid coloured circles), ancient extinct seal populations sampled for this study (opaque coloured circles: Victoria Land Coast; Tasmania; New Zealand), and ancient extinct seal populations not sampled in this study (opaque grey coloured circles). The opaque blue-grey colouring around Antarctica is the putative (see text) Last Glacial Maximum sea ice extent.

Population size at Península Valdés may have been founded within the last 6000 years (Hoelzel et al., 2001) and has been steadily increasing over the last century (Ferrari et al., 2013), contrary to historical records that suggest it was founded circa 1940s (van der Hoff et al., 2014). The South Georgia stock, and Kerguelen and Heard Island stock, have remained relatively stable since the end of sealing (*circa* 1964). In contrast, the Macquarie Island stock has been decreasing for several decades (van den Hoff et al., 2014). The causes of this decline are not entirely clear. Tracking data show that Macquarie Island seals comprise three groups that utilise different foraging zones (Hindell et al., 2017), which appear to be stable through time (Bradshaw et al., 2004): the first group forages mostly in sub-Antarctic waters, the second group largely north of the Ross Sea, and the third group (40% of tracked seals) predominantly along the Wilkes, Oates, and Victoria Land Coasts of Antarctica (Hindell et al., 2017). Increasing sea ice along these coastal regions (Hobbs et al., 2016; Hindell et al., 2017) is negatively correlated with population size at Macquarie Island, and is a likely contributing factor for the overall population size decline in this breeding stock (Hindell et al., 2017).

Phenotypic differences persist among the four stocks of southern elephant seal, likely due to low levels of (mostly paternal) gene flow. Females from Macquarie Island are smaller than females from South Georgia (Koch et al., 2019) and reach reproductive age later than in other colonies, and pup weaning weight varies between locations (Burton et al., 1997). This differentiation is thought to be result from differences in food availability (Fabiani et al., 2006; McMahon et al., 2005), and the strong philopatry displayed by females (Fabiani et al., 2003). Sexually mature females return annually to the beach on which they were born, while subadult and mature ‘bachelor’ males may disperse, sometimes over huge distances, until they establish a harem of their own (Chua et al., 2022; Fabiani et al., 2003).

Although southern elephant seals display highly philopatric behaviour, the global population has changed dramatically several times during the late Holocene. In the southern Atlantic Ocean sector, the Península Valdés (Patagonia, Argentina) stock was likely founded during the Late Holocene, and now births over 15,000 pups annually (Ferrari et al., 2013). In the Pacific sector of the Southern Ocean, Victoria Land Coast once hosted a southern elephant seal colony that was likely to have been founded around 8,000 BP, then declined to extinction beginning approximately 1,000 BP, in response to a reduction in ice-free habitat (Figure 1) (de Bruyn et al., 2009, 2014). This population expansion and eventual disappearance was probably a direct result of Holocene climate change with southwards and northwards movement, respectively, of circumpolar westerly winds and oceanic fronts and associated temperature changes (Hall et al., 2006, 2023). This now-extinct population showed close genetic affinities with seals from Macquarie Island, the extant major breeding stock also located in the southern Pacific sector (de Bruyn et al. 2009, 2014).

Another putative independent population in the southern Pacific sector is more cryptic. Extensive southern elephant seal remains in Aboriginal middens date back to at least 8,000 BP on Tasmania, Australia (Jones, 1971). Prehistoric remains of some 300 seals were excavated from West Point Midden on the north-western coast of Tasmania (Jones, 1971) (Figure 2, Figure S1, Table S1), which was occupied between approximately 2,000 and 1,000 years ago (Jones, 1971; Bryden et al., 1999). Extrapolation from the area excavated to the site as a whole suggests a minimum of 3,000 individuals, and possibly as many as 6,000 individual seals, were deposited at the West Point Midden over this time (Bryden et al., 1999). Analysis of 145 canine teeth (107 females, 38 males, 26% < 3 months old) determined age, sex, seasonal exploitation and reproductive patterns, suggesting that these seals had been procured from a nearby breeding colony (Bryden et al., 1999). There is no evidence of subsequent ongoing exploitation of elephant seals at the site after ∼1000 cal BP and Bryden et al. (1999) concluded that the nearby breeding colony became extinct at about this time. A breeding colony of southern elephant seals also existed on King Island, located between the Australian mainland and Tasmania, until the early 19^th^ Century when they were exterminated by European sealers (Bryden et al., 1999). In the absence of paleogenetic studies, however, the links between these seals and those from other populations and the four major breeding stocks are unknown.

**Figure 2.**
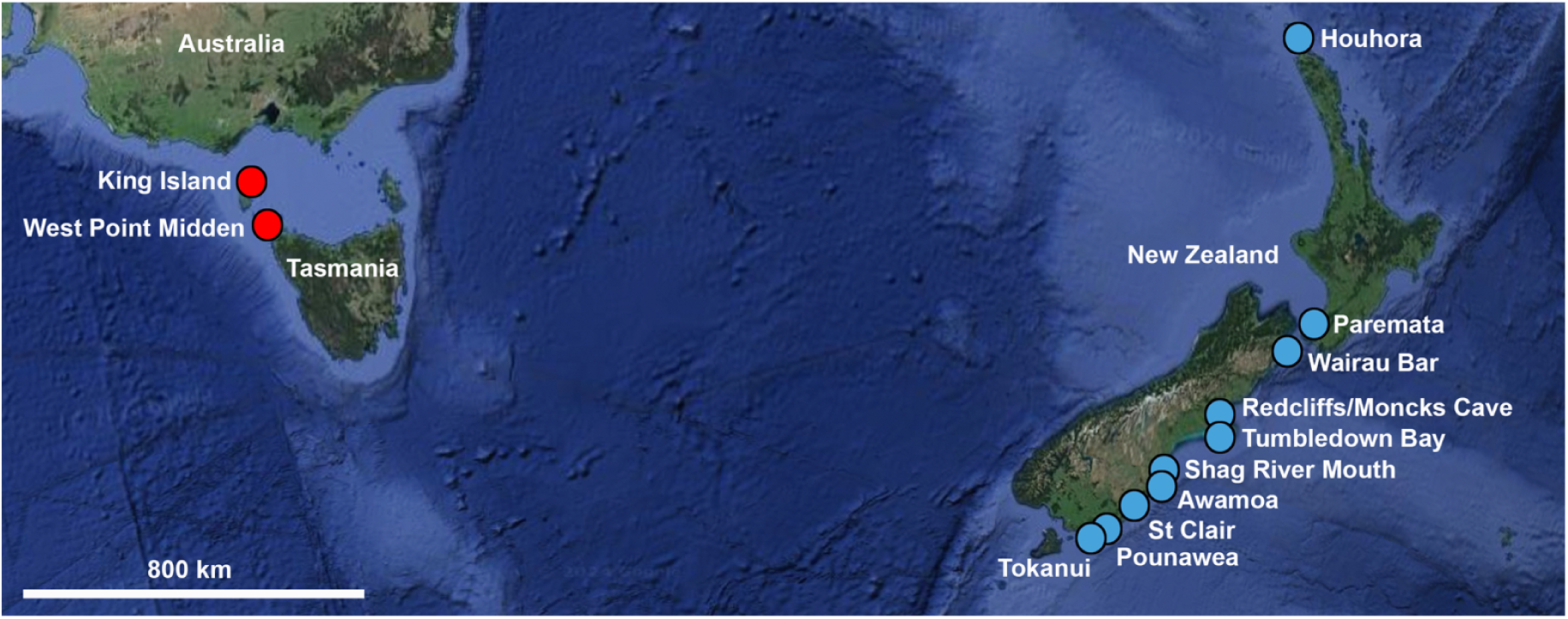
Overview of all Australian (red) and New Zealand (blue) sites from which samples of the southern elephant seal were obtained for ancient DNA analysis. Only sites with samples that yielded viable DNA are shown.

Remains of southern elephant seals have also been found in early Māori (∼1250-1450 AD) coastal middens throughout Aotearoa New Zealand in the south Pacific sector (Figure 2, Figure S2, Table S2). This species was a consistent food resource throughout New Zealand (Smith, 1985; Challis, 1995). Like the seals from Tasmania (West Point and King Island), the remains are of unknown phylogenetic/population affinity. There are fewer remains than at West Point, and their early Māori association suggests that they are at least ∼250 years younger than the youngest remains from the Tasmanian sites, i.e. <750 years BP (Smith, 1985; Challis, 1995; Bryden, 1999). There is evidence for the exploitation of southern elephant seals in New Zealand as late as 600 BP (Smith, 2013). The extant seal colony geographically closest to the New Zealand and Tasmanian mainland is located on the Campbell and Antipodes Islands, 600 and 860 km southeast of southern New Zealand, respectively (and 730 km from each other); again, little is known about these small rookeries. Macquarie Island, around 1,500 km south of Tasmania and New Zealand, forms the major extant breeding stock in this south Pacific sector.

To understand how past climate change and more recent human pressures have affected population dynamics in southern elephant seals, we sequenced mitochondrial control region and complete mitochondrial genomes (mitogenomes) from modern and ancient seals. Population genetic and phylogenetic analyses of these data sets show that extinct and extant seals from the south Pacific sector of the Southern Ocean form a distinct ‘Australasian’ lineage. Subfossil seal remains indicate that newly available habitat after the Last Glacial Maximum (LGM) allowed a major range expansion of seals. Our genetic results suggest this expansion occurred from multiple glacial refugia. However, much of this diversity was lost as climatic and human pressures have increased over the past millennium, leading to a major contraction in range size, population size, and genetic diversity of southern elephant seals.

## Results and Discussion

### Global Population Genetic Structure in the Southern Elephant Seal

We first assessed the global population genetic structure of extant and ancient southern elephant seals using various summary statistics (Table 1), AMOVA, and temporally-aware network analysis of mitochondrial control region sequences (Figure 3). These findings recapitulate previous results that split southern elephant seals into two major groups (AMOVA among groups: 35.9% variation, *p*<0.05; among populations within groups: 14.4%, *p*<0.05; within populations: 49.7%, *p*<0.05): an ‘Australasian’ lineage that includes seals from Macquarie Island and the now-extinct Victoria Land Coast population (both Pacific Ocean sector); and a lineage that includes the other extant populations (Falkland Islands, Elephant Island, Heard Island, Marion Island, South Georgia Island, King George Island, and Península Valdés) (Corrigan et al., 2016; de Bruyn et al., 2009; Chauke 2008; Fabiani et al., 2003; Slade et al., 1998; Bogdanowicz et al., 2013; Hoelzel et al., 1993). Within the latter group, finer-scale subdivision, while limited, does reflect to some degree the currently described breeding stocks: the South Georgia and Península Valdés stocks (Atlantic Ocean sector), and the Kerguelen and Heard Islands stock (Indian Ocean sector) (Figure 3). The seals in our expanded data set, which includes modern samples from the Antipodes Islands, and ancient samples from Tasmania (West Point Midden and King Island) and New Zealand, grouped with seals from Macquarie Island and Victoria Land Coast in the Australasian lineage in the Pacific Ocean sector, as expected based on geography (Figures 1–3).

**Figure 3.**
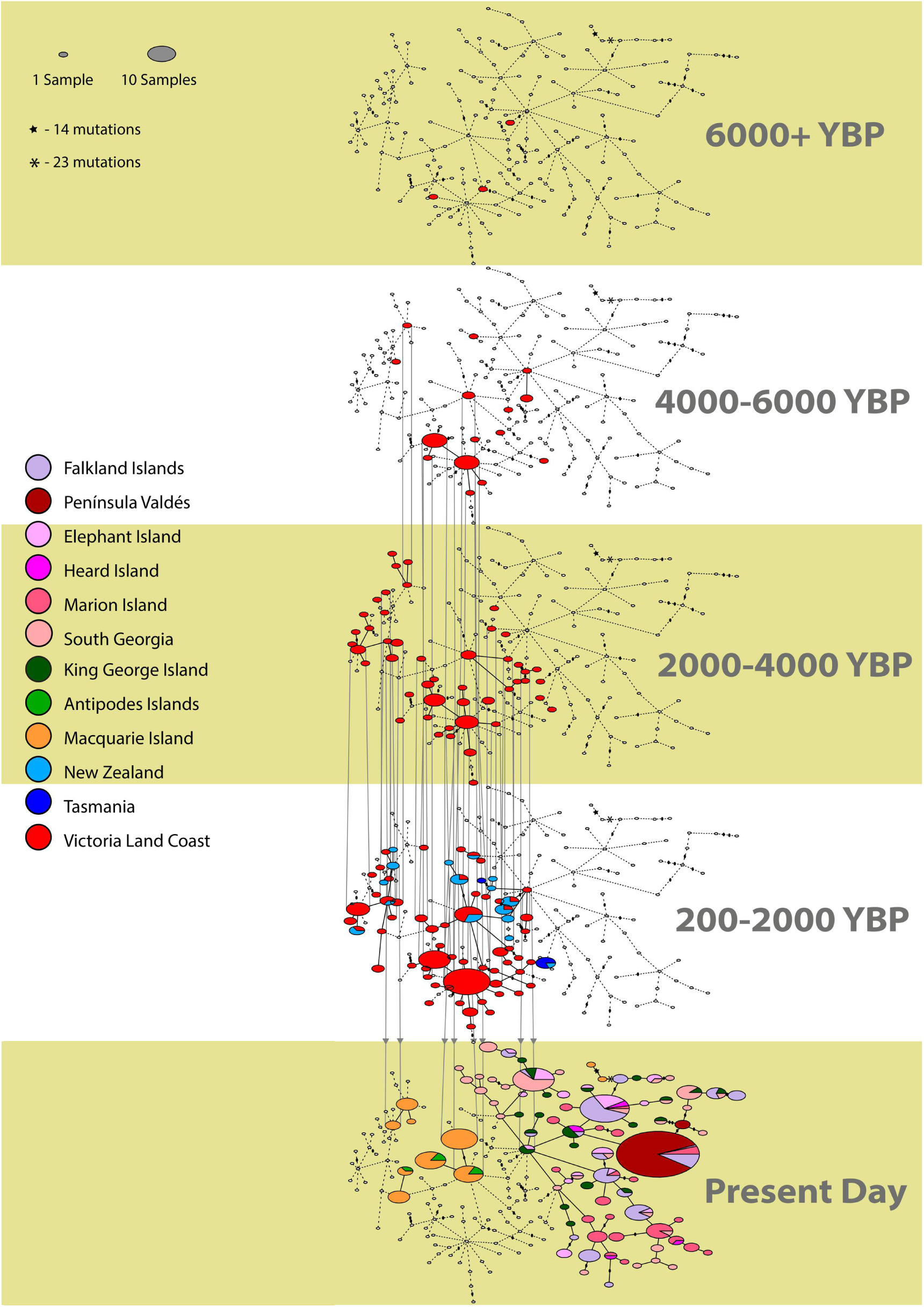
Temporally aware haplotype network showing the relationships among haplotypes for five time periods. Haplotypes are represented as ellipses, coloured by location. Ellipse size represents haplotype frequency. Lines link haplotypes separated by one mutation; dots represent additional mutations. Haplotypes absent in any particular timeframe are shown as white ellipses on that layer.

**Table 1.**
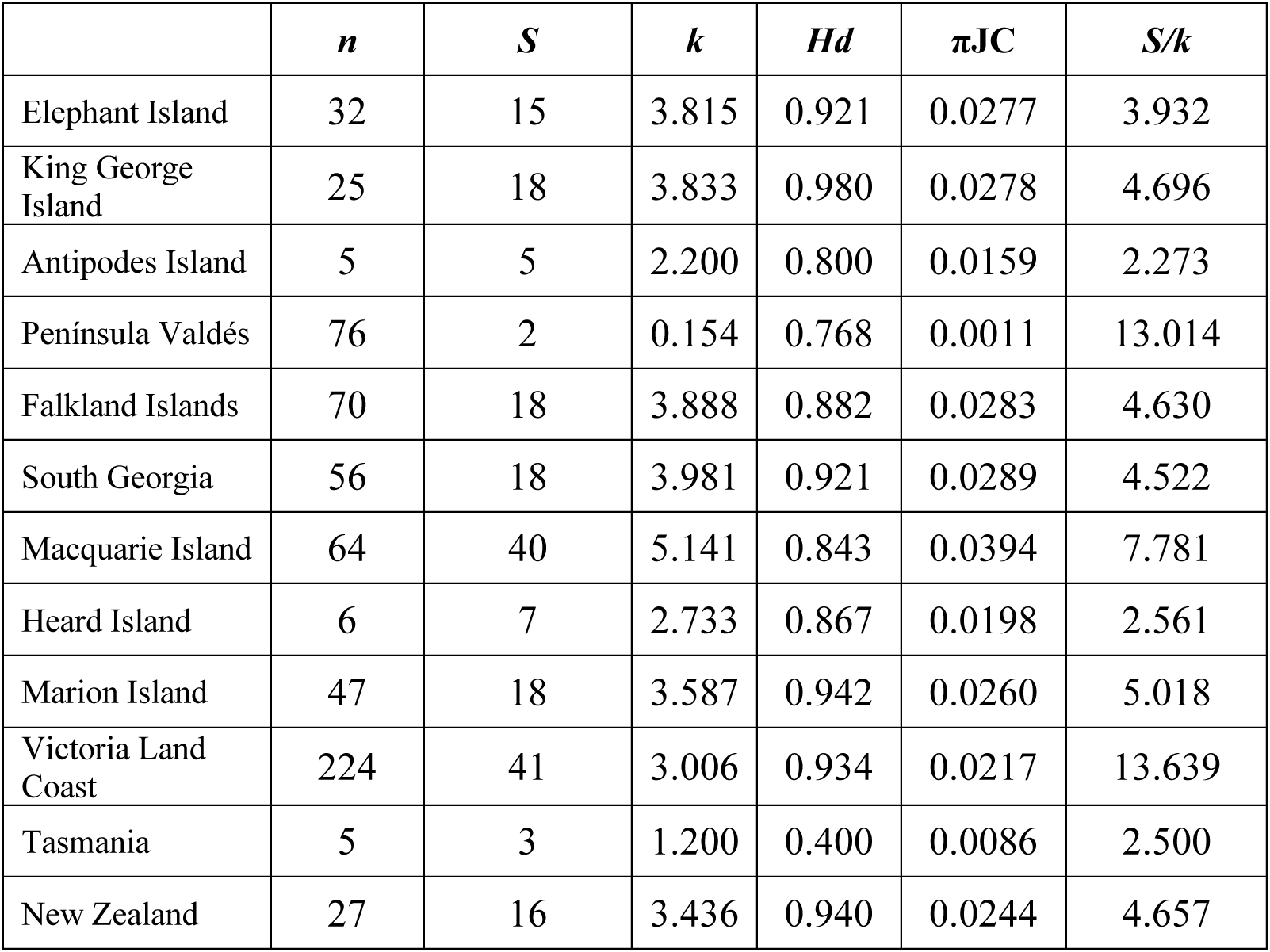
Nucleotide diversity statistics for the mitochondrial control region from populations of the southern elephant seal (*Mirounga leonina*). *n* = sample size, *S* = number of segregating sites, *k* = average number of differences, *Hd* = haplotype diversity, πJC = Jukes-Cantor corrected nucleotide diversity, *S/k* = expansion coefficient.

### Genetic Diversity and Differentiation Among Australasian Seals

To examine relationships among seals from the Australasian lineage, we conducted further population genetic analyses. The Tasmanian (including King Island) and Victoria Land Coast seals were significantly differentiated from those of all other Australasian locations, and from each other, in pairwise comparisons of control region sequences. Non-significant pairwise comparisons of differentiation were found among Macquarie Island, Antipodes Island, and New Zealand (Table S9). The Tasmanian samples appear to show extremely low genetic diversity (Table 1), with the caveat that this was the smallest sample in the data set (*n*=5, alongside Antipodes Island). In comparison, the New Zealand samples (*n*=27) showed the second-highest haplotype diversity (*Hd*) after Macquarie Island and were close to the median for all other diversity measures. The expansion coefficient (*S/k*) for the samples from Tasmania and Antipodes Island were the two lowest values found across populations. In contrast, values for the VLC (and Península Valdés, currently expanding) were an order of magnitude greater (Table 1). A caveat is that some of the populations included here are sampled across different time points, whereas other are sampled only at the present. In a population undergoing drift, sampling across a broad time range will yield a higher diversity measure in comparison to sampling at any single time point.

### Ancestry of Ancient Holocene Australasian Southern Elephant Seals

To further understand relationships among the Australasian seals, we sequenced whole mitogenomes from a subset of ancient samples from Victoria Land Coast (*n*=3), Tasmania (*n*=4), and New Zealand (*n*=3), and modern samples from Macquarie Island (*n*=12). These sequences were analysed in combination with one previously sequenced mitogenome (Accession no.: NC_008422) (Figure 4). Ancient southern elephant seals from New Zealand were most closely related to the extant Macquarie Island and extinct Victoria Land Coast populations. A Victoria Land Coast sample (3,530 BP) was the sister lineage to all other individuals in the tree, with the remaining seals forming two groups (Figure 4).

**Figure 4.**
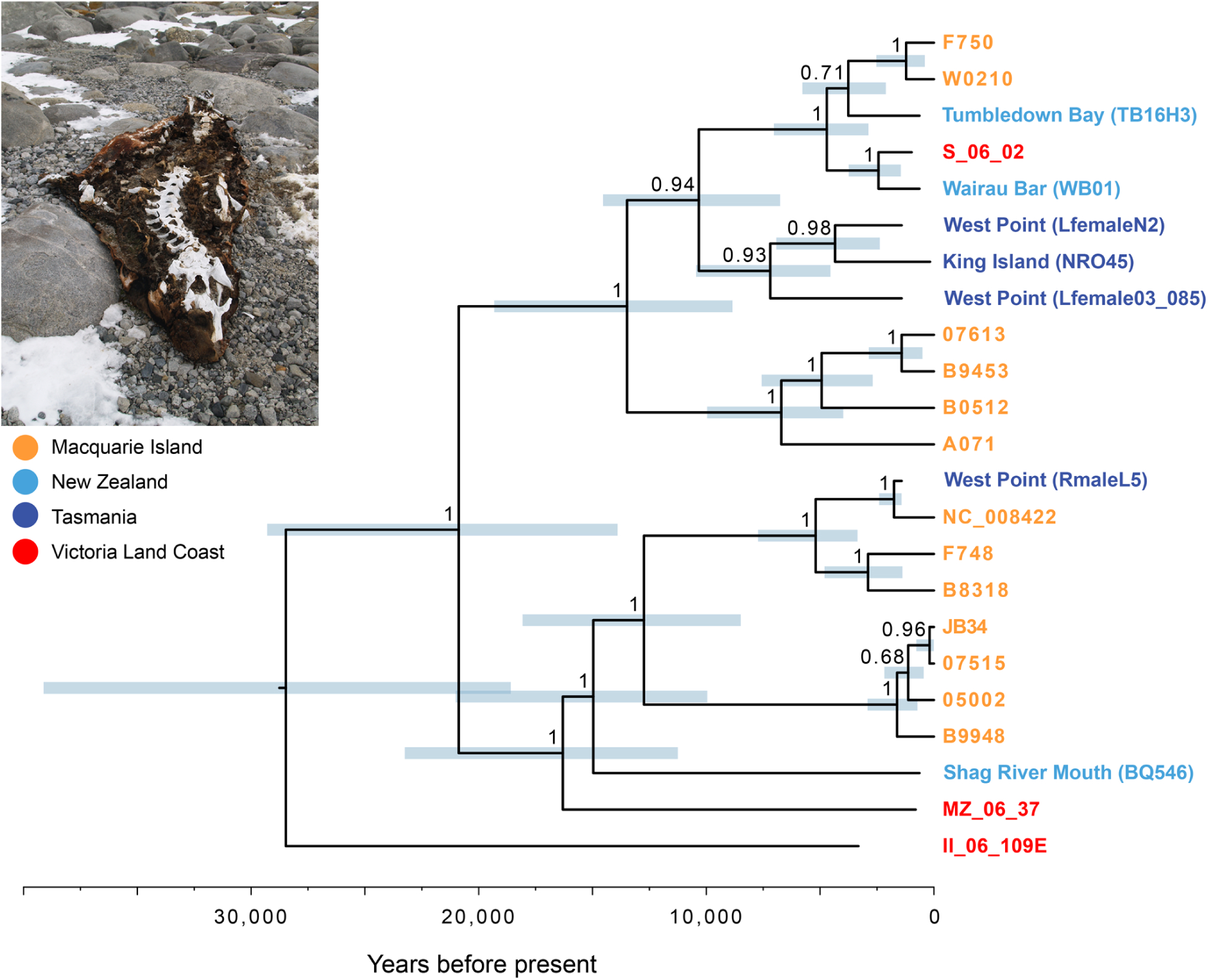
Dated phylogenetic tree from Bayesian analysis of 23 mitochondrial genomes from southern elephant seals (10 ancient & 13 modern seals). Estimates of divergence times were calibrated using the radiocarbon dates of ancient mitochondrial genomes, along with a previous estimate of the mutation rate for mitochondrial hypervariable region 1 (de Bruyn et al., 2009). Light blue bars indicate 95% credibility intervals of age estimates. Nodes are labelled with posterior probabilities. Sex of the individual is shown where known. Inset shows a mummified southern elephant seal, which previously yielded ancient DNA, from the now extinct Victoria Land Coast population (de Bruyn et al., 2009).

The temporally aware network of control region sequences showed sharing of haplotypes between ancient Victoria Land Coast seals and present-day seals from Macquarie Island and Antipodes Island, between ancient Victoria Land Coast and New Zealand seals, and a single shared haplotype between ancient New Zealand and Tasmanian seals. No haplotypes were shared between ancient Tasmanian and Victoria Land Coast seals, reflecting the significant pairwise comparisons of differentiation for these two locations. Mitogenomes further support the notion that Tasmanian seals were somewhat independent, although nested within the broader Australasian lineage, at least for the three females sampled here (Figure 4).

### Archaeological and Genetic Insights into Australasian Southern Elephant Seals

Confirmed southern elephant seal remains in an archaeological context have been found in coastal midden and occupation sites along the entire New Zealand archipelago, including the Chatham Islands, 800 km east of mainland New Zealand. These findings indicate direct human exploitation of the southern elephant seal for food and personal ornament manufacture, by East Polynesian colonists to both New Zealand and the Chatham Islands (Figure S2, Table S2). Coastal archaeological sites along Tasmania (and its associated islands in the Bass Strait) and the western, southern, and Victoria regions of the Australian mainland, and Norfolk Island (settled by Polynesians, not indigenous Australians), were also found to have southern elephant seal remains (Figure S1, Table S1). Sparse seal remains are present on Africa and South America, as well as a historic population on St Helena, off the coast of west Africa. All of these instances, excluding the Victoria Land Coast population that is presumed to have colonised and gone extinct between localised sea-ice retreat and expansion events, 8,000–1,000 BP (de Bruyn et al., 2009), occur north of the estimated Last Glacial Maximum winter sea-ice extent (from Fraser et al., 2009; Gersonde et al., 2005; Rawlence et al., 2022; Trucchi et al., 2014).

The majority of documented prehistoric remains of southern elephant seals are younger than the Last Glacial Maximum (21,000–18,000 BP, Tables S1–S3), the only exceptions being rare moulted skin from the Victoria Land Coast of Antarctica immediately prior to the LGM (Hall et al., 2023), and late Pleistocene fossils from South America (Valenzuela-Toro et al., 2015) and New Zealand (Boessenecker and Churchill, 2016), supporting the hypothesis that these areas acted as glacial refugia. Fossils older than the Pleistocene-Holocene boundary ∼10Kya, and indeed the LGM, are comparatively rare and less likely to be found. The comparatively young age of many of the prehistoric specimens is likely to be a product of older coastal breeding sites being inundated at the end of the LGM by rising sea levels until ∼6,000 BP (Lambeck and Nakada, 1990). When Antarctic sea ice advanced during the LGM, it displaced local pagophobic (ice-intolerant) species including the southern elephant seal, causing their ranges to shift northward. When the ice started to recede after the LGM, new locations were open for exploitation. This is evidenced by the postglacial expansions inferred for many different taxa around Antarctica (Fraser et al., 2009; Fraser et al., 2012; Gonzalez-Wevar et al., 2018; Nikula et al., 2010; Rawlence et al., 2022). At the end of the LGM ∼17.5 Kya, there was rapid warming of southern mid- to high-latitudes facilitated by southward movement of circumpolar westerly winds and ocean fronts, which would have led to rapid southward range expansions into the sub-Antarctic of marine taxa up to the limit of the retreating sea ice, with Antarctica lagging behind (13 Kya for the Ross Sea and 8 Kya for Victoria Land Coast).

Our analyses strongly suggest that, at the end of the LGM, the southern elephant seal had a markedly different breeding range from what is seen today, reaching well into the current temperate zone before expanding deep into the sub-Antarctic and Antarctica during the Holocene, before contracting into the sub-Antarctic due to sealing pressures (Bryden et al., 1999; Thatje et al., 2008). New Zealand, Tasmania, and some of the surrounding sub-Antarctic islands, such as Campbell Island, the Auckland Islands, and potentially even the Chatham Islands, acted as important ice-free refugia for the Australasian southern elephant seal breeding stock when the current stronghold of Macquarie Island was likely impacted by sea ice during the LGM (Fraser et al., 2009; Gersonde et al., 2005). The southern reaches of Africa and South America may also have acted as ice-free refugia for populations found now on islands that would have been displaced by encroaching sea ice, such as King George Island, Elephant Island, and South Georgia (Fraser et al., 2009; Hall, 2004; Rainsley et al., 2019; Rawlence et al., 2022; Rawlence et al., 2024, but see Gersonde et al., 2005; Scott and Turnbull, 2019). This hypothesis is potentially supported by the presence of late Pleistocene southern elephant seal remains on these continents (Avery and Klein, 2011; Valenzuela-Toro et al., 2015). However, the scarcity of southern elephant seal remains in these areas (see Valenzuela-Toro et al., 2015) is again probably due to sea level changes causing coastal sites to be lost, making their status as breeding populations impossible to verify at this stage.

Temporal comparisons revealed that many ancient samples of southern elephant seals across New Zealand had mitochondrial haplotypes identical to modern samples from Macquarie and Antipodes islands (Fig. 3). These findings suggest that the prehistoric New Zealand population was genetically connected to other Australasian lineage populations (*cf*. *Phocarctos* sea lions; Collins et al., 2013; Collins et al., 2014; Rawlence et al., 2016). While genetic evidence remains inconclusive about the status of New Zealand as a breeding site, archaeological evidence is somewhat contradictory. The ubiquity and sudden disappearance of elephant seals shortly after human settlement of New Zealand, and their lack of replacement (*cf*. Boessenkool et al., 2009; Collins et al., 2014; Rawlence et al., 2015; Waters et al., 2017) prior to European exploitation of sub-Antarctic populations in the late 18^th^ Century, show a regionally consistent pattern of decline over the entire archipelago (Nagaoka, 2000; Smith, 1989; Smith, 1996; Smith, 2011).

All archaeological sites with southern elephant seal in New Zealand have been dated (either through radiocarbon dating or associated cultural assemblage) to ∼1250-1450 AD (Table S2), indicating a sudden disappearance of southern elephant seals from the region coincident with the extinction (e.g., moa; Holdaway et al., 2014; Perry et al. 2014; *Phocarctos* sea lions; Collins et al., 2013; Rawlence et al. 2016; Waters et al., 2017; seabirds; Boessenkool et al., 2009; Rawlence et al., 2015; Rawlence et al., 2017) and severe range-contractions and population bottlenecks (e.g., *Leucocarbo* shags; Rawlence et al., 2015; *Arctocephalus* furseals; Ling, 1999b; Ling, 2000; Salis et al., 2016) seen in terrestrial and marine megafauna. We do not see any evidence for recolonisation events (e.g., Boessenkool et al., 2009; Collins et al., 2014). This rapid decline suggests a direct extirpation of mainland breeding populations. Similarly, there is compelling archaeological evidence of former southern elephant seal breeding populations, numbering hundreds or thousands of individuals, including females and pups, on the Tasmanian mainland and King Island (Bryden et al., 1999). On King Island there is additional historical evidence of a small breeding colony that was exploited to the point of extinction by European sealers (Ling, 1999a).

If the New Zealand mainland had been host only to occasional vagrants or moulting individuals from nearby populations (such as those on Campbell Island or Antipodes Islands), we would expect a reasonably constant presence/proportion of southern elephant seals throughout the pre-European and historic archaeological record. The consistent replacement from nearby, unexploited populations would have been similar to the pattern seen in leopard seals (*Hydrurga leptonyx*) (Anderson, 1983; Smith, 2011; Smith, 2013), *Phocarctos* sealions, *Megadyptes* and *Eudyptula* penguins, and *Cygnus* swans (Grosser et al., 2016; Rawlence et al., 2017; Waters et al., 2017). The shared haplotypes among New Zealand, Antipodes Island, and Macquarie Island may be due to the former two (along with the Campbell Islands) being sea ice-free refuges during the LGM when many of the sub-Antarctic islands south of the winter LGM sea ice extent (Macquarie Island potentially included) were covered and/or surrounded in ice (Fraser et al., 2009; Hall, 2004; Rainsley et al., 2019; Rawlence et al., 2022; Rawlence et al., 2024, but see Gersonde et al., 2005; Scott and Turnbull, 2019). Population range expansion from the New Zealand region to Macquarie Island likely occurred shortly after the sea ice receded, similar to the colonisation of Victoria Land Coast from Macquarie Island inferred by de Bruyn et al. (2009).

When mitochondrial control region sequences for individuals from West Point Midden in Tasmania were compared with other sequences in the global network, they formed a distinct group; female mitogenomes form a similarly distinctive subclade (Fig. 4). This inferred presence of a distinct Tasmanian breeding population is further supported by archaeological evidence, where multiple sites with abundant remains, including those of pups, are present along Tasmania’s coastline for extended periods (Jones, 1971; Bryden et al., 1999). Due to Tasmania’s early colonisation by humans by at least 43,000 BP (Gillespie et al., 2012), any coastal geomorphological changes that may have occurred with changes in sea levels must also be taken into account. Rising sea levels since the LGM may have contributed to many potential older archaeological southern elephant seal butchery sites now being submerged. Tasmanian sea levels began to stabilise ∼6,000 BP (Lambeck and Nakada 1990), which coincides with the dates of many of the elephant seal butcheries found on the Tasmanian mainland (Jones, 1971; Bryden et al., 1999). The perseverance of elephant seal remains in the mainland Tasmanian archaeological record from the mid- to late Holocene (from at least 8,000 BP; Jones 1971) might be attributable to, in part, recolonisation from refuge populations inhabiting offshore islands, such as King Island, which would have been isolated from the mainland by 6,000 BP. The presence of elephant seal remains on three distinct islands in this region by the Late Holocene (Figure S1, Table S1), including the breeding population exploited by European sealers on King Island (Ling, 1999a; Ling, 2002), lends credence to the hypothesis that there was at least one healthy offshore population acting as a reservoir for the exploited mainland population(s) during the mid- to late Holocene; ensuring the sustainability of Aboriginal subsistence hunting.

The results of this study contribute to a growing body of evidence that demonstrates how polar and sub-polar species have historically responded to significant environmental changes. For example, during the Last Glacial Maximum (LGM), many species adapted by shifting their ranges to available refugia, as evidenced by genetic data from kelp-dwelling crustaceans and other marine organisms (Nikula et al., 2010; Fraser et al., 2009). These adaptive responses, while showcasing resilience, also highlight the limits of such shifts when combined with anthropogenic pressures. Modern parallels can be drawn from current observations, such as declining populations of polar bears due to shrinking ice habitats and reduced hunting grounds, or changes in penguin breeding success as sea ice patterns alter access to feeding areas (Hindell et al., 2017; Vianna et al., 2020). By understanding the past resilience and vulnerabilities of polar and sub-polar species, conservationists can develop more targeted approaches to support species facing the dual challenges of warming temperatures and human-induced habitat disruption (Hoegh-Guldberg & Bruno, 2010; Rawlence et al., 2022).

## Conclusion

In the Southern Ocean, a region currently undergoing rapid environmental change, prehistoric climate change (associated with the LGM, Pleistocene-Holocene transition, and regional/local changes in sea ice extent) and human pressures over the past few thousand years in the Pacific sector substantially altered the distribution of southern elephant seals over short evolutionary timescales. Genetic data, and historic, archaeological, and subfossil remains indicate that a post-LGM expansion from middle latitude refuges on Australia, New Zealand and perhaps other locations, and sealing-induced contraction from multiple groups of humans have shaped Australasian populations of the southern elephant seal. Analyses of these diverse datasets illuminates the extent of ‘dark intraspecific extinction’ in a large, well-studied mammalian system; that is, a significant loss of within-species diversity (Figs. 3 & 4), and a major reduction in geographic range (Figs. 1 & 2), due to human and environmental pressures during the late Holocene.

## Materials and Methods

### HVR1 Sequence Analysis

Partial HVR1 sequences were successfully amplified from modern and ancient samples. A data set with representative sequences from various populations was constructed. Genetic diversity metrics were calculated, and FST values were used to measure genetic differentiation. AMOVAs were performed to infer geographic structure. A time-sensitive haplotype network was constructed (SI Appendix).

### Mitochondrial Genome Assembly

DNA extraction was performed on selected ancient samples for mitochondrial genome assembly. Libraries were prepared, quantified, and enriched using hybridization-capture with bait from a closely related species. PCR amplification and pooling of samples were performed, and sequenced using the Illumina MiSeq platform (SI Appendix).

### Data Processing

Sequence data were processed using specific scripts to trim adapters, merge overlapping read pairs, and filter low-quality reads. The processed reads were mapped to a reference mitochondrial genome using BWA and filtered for quality and duplicates. Consensus sequences were computed, and patterns of DNA damage were assessed (SI Appendix).

### Phylogenetic Analysis of Mitogenome Sequences

Newly generated mitogenome sequences were aligned with global southern elephant seal sequences.

Post-mortem damage was assessed using MapDamage (SI Appendix). We also used Bayesian analysis in BEAST v1.8.2 (Suchard et al. 2018) to compare three models of post-mortem sequence damage: no damage, age-dependent damage, and age-independent damage (Ho et al., 2007; Rambaut et al., 2009; Ho, 2012). Models were compared using marginal likelihoods estimated by stepping-stone sampling (Xie et al., 2010). Posterior distributions of parameters were estimated using Markov Chain Monte Carlo sampling over 75 million steps, with samples drawn every 10^4^ steps.

To infer a dated phylogenetic tree, we performed a Bayesian molecular dating analysis in BEAST. We used a skyride coalescent tree prior (for species with a dynamic population history) and a strict molecular clock, selected by comparison of marginal likelihoods calculated by stepping-stone sampling. Estimates of node times were calibrated using the sample dates for the ancient Tasmanian, New Zealand, and Victoria Land Coast sequences. We also fixed the mutation rate of HVR1 based on an estimate from a previous study of the southern elephant seal (de Bruyn et al., 2009). Posterior distributions of parameters were estimated using Markov Chain Monte Carlo sampling over 75 million steps, with samples drawn every 10^4^ steps. We ran the analysis in duplicate to check for convergence, and combined the samples after discarding the first 10% as burn-in. Sufficient sampling was confirmed by inspecting the samples in Tracer v1.7.1 (Rambaut et al., 2018).

## Acknowledgments

We acknowledge that Māori, the indigenous people of Aotearoa New Zealand, have kaitiaki (guardianship) over the organisms from their rohe (tribal area). BH acknowledges support from the Office of Polar Programs of the US National Science Foundation. PF is a recipient of an ARC Future Fellowship (FTFT200100464).

## References

Allentoft, M.E., Heller, R., Oskam, C.L., Lorenzen, E.D., Hale, M.L., Gilbert, M.T.P., Jacomb, C., Holdaway R.N., Bunce, M. (2014) Extinct New Zealand megafauna were not in decline before human colonization. Proceedings of the National Academy of Sciences of the United States of America, 111, 4922–4927.

Anderson, A. (1983) Faunal depletion and subsistence change in the early prehistory of southern New Zealand. Archaeology in Oceania, 18, 1–10.

Avery, G. and Klein, R.G. (2011) Review of fossil phocid and otariid seals from the southern and western coasts of South Africa. Transactions of the Royal Society of South Africa, 66, 14–24.

Boessenecker, R.W., Churchill, M. (2016) The origin of elephant seals: implications of a fragmentary late Pliocene seal (Phocidae: Miroungini) from New Zealand. New Zealand Journal of Geology and Geophysics, 59, 544–550.

Boessenkool, S., Austin, J.J., Worthy, T.H., Scofield, P., Cooper, A., Seddon, P.J., Waters, J.M. (2009) Relict or colonizer? Extinction and range expansion of penguins in southern New Zealand. Proceedings of the Royal Society B, 276, 815–821.

Bogdanowicz, W., Pilot, M., Gajewska, M., Suchecka, E. & Golachowski, M. (2013) Genetic diversity in a moulting colony of southern elephant seals in comparison with breeding colonies. Marine Ecology Progress Series, 478, 287–300.

Bradshaw, C.J.A., Hindell, M.A., Sumner, M.D., Michael, K.J. (2004) Loyalty pays: potential life history consequences of fidelity to marine foraging regions by southern elephant seals. Animal Behaviour, 68, 1349–1360.

Bryden, M. M., O’Connor, S. & Jones, R. (1999) Archaeological evidence for the extinction of a breeding population of elephant seals in Tasmania in prehistoric times. International Journal of Osteoarchaeology, 9, 430–437.

Burton, H.R., Arnbom, T., Boyd, I.L., Bester, M., Vergani, D., Wilkinson, I. (1997) Significant differences in weaning mass of southern elephant seals from five sub-Antarctic islands in relation to population declines. In: Antarctic Communities: Species, Structure and Survival (eds Battaglia, B., Valencia, J., Walton, D.). Cambridge University Press. pp. 335–338.

Challis, A. J. (1995) *Ka pakihi whakatekateka o waitaha: the archaeology of Canterbury in Maori times*: Department of Conservation Wellington.

Chauke, L.F. (2008) Genetic variation and population structure of southern elephant seals *Mirounga leonina* from Marion Island. The University of Pretoria. PhD Thesis.

Chua, M., Ho, S.Y.W., McMahon, C.R., Jonsen, I.D., de Bruyn, M. (2022) Movements of southern elephant seals (Mirounga leonine) from Davis Base, Antarctica: combining population genetics and tracking data. Polar Biology, 45, 1163–1174.

Cole, T.L., Dutoit, L., Dussex, N., Hart, T., Alexander, A., Younger, J.L., Clucas, G.V., Frugone, M.J., Cherel, Y., Cuthbert, R., et al. (2019) Receding ice drove parallel expansions in Southern Ocean penguins. PNAS, 116, 26990–26996.

Collins, C.J., Rawlence, N.J., Worthy, T.H., Scofield, R.P., Tennyson, A.J.D., Smith, I., Knapp, M., Waters, J.M. (2013) Pre-human New Zealand sea lion (*Phocarctos hookeri*) rookeries on mainland New Zealand. Journal of the Royal Society of New Zealand, 44, 1–16.

Collins, C.J., Rawlence. N.J., Prost. S., Anderson. C., Knapp. M., Scofield. R.P., Robertson. B.C., Smith. I., Matisoo-Smith. E.A., Chilvers. B.L., Waters. J.M. (2014) Extinction and recolonisation of coastal megafauna following human arrival in New Zealand. Proceedings of the Royal Society B, 281, 20140097.

Corrigan, L.J., Fabiani, A., Chauke, L., McMahon, C.R., de Bruyn, M., Bester, M.N., Bastos, A., Campagna, C., Muelbert, M.M., Hoelzel, A.R. (2016) Population differentiation in the context of Holocene climate change for a migratory marine species, the southern elephant seal. Journal of Evolutionary Biology, 29, 1667–1679.

de Bruyn, M., Hall, B.L., Chauke, L.F., Baroni, C., Koch, P.L., Hoelzel, A.R. (2009). Rapid response of a marine mammal species to Holocene climate and habitat change. PLOS Genetics, 5, e1000554.

de Bruyn, M., Pinsky, M., Hall, B., Koch, P., Baroni, C., Hoelzel, A.R. (2014). Rapid increase in southern elephant seal variation after a founder event. Proceedings of the Royal Society B, 281, 20133078.

Fabiani, A., Galimberti, F., Sanvito, S., Hoelzel, A.R. (2006). Relatedness and site fidelity at the southern elephant seal, *Mirounga leonina*, breeding colony in the Falkland Islands. Animal Behaviour, 72, 617–626.

Fabiani, A., Hoelzel, A.R., Galimberti, F., Muelbert, M.M.C. (2003). Long-range paternal gene flow in the southern elephant seal. Science, 299, 676.

Ferrari, M., Campagna, C., Condit, R. & Lewis, M. (2013). The founding of a southern elephant seal colony. Marine Mammal Science, 29, 407–423.

Fraser, C.I., Nikula, R., Ruzzante, D.E., Waters, J.M. (2012). Poleward bound: biological impacts of Southern Hemisphere glaciation. Trends in Ecology and Evolution, 27, 462– 471.

Fraser, C.I., Nikula, R., Spencer, H.G., Waters, J.M. (2009). Kelp genes reveal effects of subantarctic sea ice during the Last Glacial Maximum. Proceedings of the National Academy of Sciences of the United States of America, 106, 3249–3253.

Gales, N.J., Adams, M., Burton, H.R. (1989). Genetic relatedness of two populations of southern elephant seal (*Mirounga leonina*). Marine Mammal Science, 5, 57–67.

Gersonde, R., Crosta, X., Abelmann, A., Armand, L. (2005). Sea-surface temperature and sea ice distribution of the Southern Ocean at the EPILOG Last Glacial Maximum–a circum-Antarctic view based on siliceous microfossil records. Quaternary Science Reviews, 24, 869–896.

Gillespie, R., Camens, A.B., Worthy, T.H., Rawlence, N.J., Reid, C., Bertuch, F., Levchenko, V., Cooper, A. (2012). Man and megafauna in Tasmania: Closing the gap. Quaternary Science Reviews, 37, 38–47.

Gonzalez-Wevar, C.A., Segovia, N.I., Rosenfeld, S., Ojeda, J., Hune, M., Naretto, J., Saucede, T., Brickle, P., Morley, S., Feral, J-P., Spencer, H.G., Poulin, E. (2018). Unexpected absence of island endemics: long-distance dispersal in higher latitude sub-Antarctic *Siphonaria* (Gastropoda: Euthyneura) species. Journal of Biogeography, 45, 874–884.

Grosser S., Rawlence, N.J., Anderson, C.N.K., Smith, I.W.G., Scofield, R.P., Waters, J.M. (2016). Invader or resident? Ancient-DNA reveals rapid species turnover in New Zealand little penguins. Proceedings of the Royal Society B, 283, 20152879.

Hall, K. (2004). Quaternary Glaciations Extent and Chronology - Part III: South America, Asia, Africa, Australasia, Antarctica, Elsevier.

Hall, B.L., Hoelzel, A.R., Baroni, C., Denton, G.H., Le Boeuf, B.J., Overturf, B., Topf, A.L. (2006). Holocene elephant seal distribution implies warmer-than-present climate in the Ross Sea. Proceedings of the National Academy of Sciences of the United States of America, 103, 10213–10217.

Hall, B.L., Koch, P.L., Baroni, C., Salvatore, M.C., Hoelzel, A.R., de Bruyn, M., Welch, A.J. (2023). Widespread southern elephant seal occupation of the Victoria Land coast implies a warmer-than-present Ross Sea in the mid-to-late Holocene. Quaternary Science Reviews, 303, 107991.

Hindell, M.A., Sumner, M., Bestley, S., Wotherspoon, S., Harcourt, R.G., Lea, M-A., Alderman, R., McMahon, C.R. (2017). Decadal changes in habitat characteristics influence population trajectories of southern elephant seals. Global Change Biology, 23, 5136–5150.

Hobbs, W.R., Massom, R., Stammerjohn, S., Reid, P., Williams, G., Meier, W. (2016). A review of recent changes in Southern Ocean sea ice, their drivers and forcings. Global and Planetary Change, 143, 228–250.

Hoegh-Guldberg, O., Bruno, J.F. (2010). The impact of climate change on the world’s marine ecosystems. Science, 328, 1523–1528.

Hoelzel, A.R., Campagna, C., Arnbom, T. (2001). Genetic and morphological differentiation between island and mainland southern elephant seal populations. Proceedings of the Royal Society of London Series B: Biological Sciences, 268,325–332.

Hoelzel, A., Halley, J., O’Brien, S. J., Campagna, C., Arnborm, T., Le Boeuf, B., Ralls, K., Dover, G. (1993). Elephant seal genetic variation and the use of simulation models to investigate historical population bottlenecks. Journal of Heredity, 84, 443–449.

Holdaway, R.N. (2000). Rapid extinction of the moas (Aves: Dinornithiformes): Model, test, and implications. Science, 287, 2250–2254.

Holdaway, R.N., Allentoft, M.E., Jacomb, C., Oskam, C.L., Beavan, N.R., Bunce, M. (2014). An extremely low-density human population exterminated New Zealand moa. Nature Communications, 5, 5436.

Jones, R. (1971). Rocky Cape and the problem of the Tasmanians. University of Sydney. PhD Thesis.

Koch, P.L., Hall, B.L., de Bruyn, M., Hoelzel, A.R., Baroni, C., Salvatore, M.C. (2019) Mummified and skeletal southern elephant seals (*Mirounga leonina*) from the Victoria Land Coast, Ross Sea, Antarctica. Marine Mammal Science, 35, 934–956.

Lambeck, K., Nakada, M. (1990). Late Pleistocene and Holocene sea-level change along the Australian coast. Palaeogeography, Palaeoclimatology, Palaeoecology, 89, 143–176.

Ling, J. K. (1999a). Elephant seal oil cargoes from King Island, Bass Strait, 1802-1819: with estimates of numbers killed and size of the original population. Papers and Proceedings of the Royal Society of Tasmania, 133, 51–56.

Ling, J.K. (1999b). Exploitation of fur seals and sea lions from Australian, New Zealand and adjacent subantarctic islands during the eighteenth, nineteenth and twentieth centuries. Australian Zoologist, 31, 323–350.

Ling, J.K., (2002). Impact of colonial sealing on seal stocks around Australia, New Zealand and subantarctic islands between 150 and 170 degrees East. Australian Mammalogy, 24, 117–126.

McMahon, C.R., Bester, M.N., Burton, H.R., Hindell, M.A., Bradshaw, C.J.A. (2005). Population status, trends and a re-examination of the hypotheses explaining the recent declines of the southern elephant seal *Mirounga leonina*. Mammal Review, 35, 82–100.

Nagaoka, L. (2000). Resource Depression, Extinction, and Subsistence Change in Prehistoric Southern New Zealand. University of Washington.

Nikula, R., Fraser, C.I., Spencer, H.G., Waters, J.M. (2010). Circumpolar dispersal by rafting in two subantarctic kelp-dwelling crustaceans. Marine Ecology Progress Series, 405, 221–230.

Perry, G.L.W., Wheeler, A.B., Wood, J.R., Wilmshurst, J.M. (2014). A high-precision chronology for the rapid extinction of New Zealand moa (Aves, Dinornithiformes). Quaternary Science Reviews, 105, 126–135.

Rainsley, E., Turney, C.S.M., Golledge, N.R., Wilmshurst, J.M., McGlone, M.S., Hogg, A.G., Li, B., Thomas, Z.A., Roberts, R., Jones, R.T., Palmer, J.G., Flett, V., de Wet, G., Hutchinson, D.K., Lipson, M.J., Fenwick, P., Hines, B.R., Binetti, U., Fogwill, C.J. (2019). Pleistocene glacial history of the New Zealand subantarctic islands. Climate of the Past, 15, 423–448.

Rawlence, N.J., Collins, C.J., Anderson, C.N.K., Maxwell, J.J., Smith, I.W.G., Robertson, B.C., Knapp, M., Horsburgh, K.A., Stanton, J.A.L., Scofield, R.P., Tennyson, A.J.D., Matisoo-Smith, E.A., Waters, J.M. (2016). Human-mediated extirpation of the unique Chatham Islands sea lion and implications for the conservation management of remaining New Zealand sea lion populations. Molecular Ecology, 25, 3950–3961.

Rawlence, N.J., Kardamaki, A., Easton, L.J., Tennyson, A.J.D., Scofield, R.P., Waters, J.M. (2017). Ancient DNA and morphometrics analysis reveal extinction and replacement of New Zealand’s unique black swans. Proceedings of the Royal Society B, 284, 20170876.

Rawlence, N.J., Kennedy, M., Anderson, C.N.K., Prost, S., Till, C.E., Smith, I.W.G., Scofield, R.P., Tennyson, J.D., Hamel, J. (2015). Geographically contrasting biodiversity reductions in a widespread New Zealand seabird. Molecular Ecology, 24, 4605–4616.

Rawlence, N.J., Perry, G.L.W., Smith, I.W.G., Scofield, R.P., Tennyson, A.J.D., Matisoo-Smith, E.A., Boessenkool, S., Austin, J.J., Waters, J.M. (2015). Radiocarbon-dating and ancient DNA reveal rapid replacement of extinct prehistoric penguins. Quaternary Science Reviews, 112, 59–65.

Rawlence, N.J., Salis, A.T., Spencer, H.G., Waters, J.M., Scarsbrook, L., Mitchell, K.J., Phillips, R.A., Calderon, L., Cook, T.R., Bost, C.A., Dutoit, L., King, T.M., Masello, J.F., Nupen, J.F., Quillfeldt, P., Ratcliffe, N., Ryan, P.G., Till, C.E., Kennedy, M. (2022). Rapid radiation of Southern Ocean shags in response to receding sea ice. Journal of Biogeography, 49, 942–953.

Rawlence, N.J., Till, C.E., Easton, L.J., Spencer, H.G., Schuckard, R., Melville, D.S., Scofield, R.P., Tennyson, A.J.D., Rayner, M.J., Waters, J.M., Kennedy, M. (2017). Speciation, range contraction and extinction in the endemic New Zealand King Shag complex. Molecular Phylogenetics and Evolution, 115, 197–209.

Rawlence, N.J., Verry, A.J.F., Cole, T.L., Shepherd, L.D., Tennyson, A.J.D., Williams, M., Wood, J.R., Mitchel, K.J. (2024). Ancient mitogenomes reveal evidence for the late Miocene dispersal of mergansers to the Southern Hemisphere. Zoological Journal of the Linnean Society, doi10.1093/zoolinnean/zlae040.

Salis, A.T., Easton, L.J., Robertson, B.C., Gemmell, N., Smith, I.W.G., Weisler, M.I., Waters, J.M., Rawlence, N.J. (2016). Myth or relict: Does ancient-DNA detect the enigmatic Upland Seal? Molecular Phylogenetics and Evolution, 97, 101–106.

Scott, J.M., Turnbull, I.M. (2019). Geology of New Zealand’s sub-Antarctic islands. New Zealand Journal of Geology and Geophysics, 62, 291–317.

Slade, R. W., Moritz, C., Hoelzel, A. R., Burton, H. R. (1998). Molecular population genetics of the southern elephant seal *Mirounga leonina*. Genetics, 149, 1945–1957.

Smith, I. W. (1985). Sea mammal hunting and prehistoric subsistence in New Zealand. University of Otago.

Smith, I.W.G. (1989). Maori Impact on the marine megafauna: Pre-European distribution of New Zealand sea mammals. In Saying So Doesn’t Make It So. Papers in Honour of B. Foss Leach. D.G. Sutton ed. Dunedin, New Zealand Archaeological Association Monograph 17, 76–108.

Smith, I.W.G. (1996). Historical documents, archaeology and 18th century seal hunting in New Zealand. In *Oceanic Culture History: Essays in Honour of Roger Green.* J.M. Davidson, G. Irwin, B.F. Leach, A. Pawley and D. Brown eds. Dunedin, New Zealand Journal of Archaeology Special Publication.

Smith, I.W.G. (2011). Estimating the magnitude of pre-European Maori marine harvest in two New Zealand study areas. New Zealand Aquatic Environment and Biodiversity Report No. 82.

Smith, I.W.G. (2013). Pre-European Maori exploitation of marine resources in two New Zealand case study areas: species range and temporal change. Journal of the Royal Society of New Zealand, 43, 1–37.

Thatje, S., Hillenbrand, C.-D., Mackensen, A., Larter, R. (2008). Life hung by a thread: Endurance of Antarctic fauna in glacial periods. Ecology, 89, 682–692.

Trucchi, E., Gratton, P., Whittington, J.D., Cristofari, R., Maho, Y.L., Stenseth, N.C., Bohec, C.E. (2014). King penguin demography since the last glaciation inferred from genome-wide data. Proceedings of the Royal Society of London. Series B: Biological Sciences, 281, 20140528.

Valenzuela-Toro, A.M., Gutstein, C.S., Suárez, M.E., Otero, R., Pyenson, N.D. (2015). Elephant seal (*Mirounga* sp.) from the Pleistocene of the Antofagasta Region, northern Chile. Journal of Vertebrate Paleontology, 35, e918883.

van den Hoff, J., McMahon, C.R., Simpkins, G.R., Hindell, M.A., Alderman, R., Burton, H.R. (2014). Bottom-up regulation of a pole-ward migratory predator population. Proceedings of the Royal Society of London. Series B: Biological Sciences, 281, 20132842.

Vianna, J.A., Fernandes, F.A.N., Frugone, M.J., Figueiro, H.V., Pertierra, L.R., Noll, D., Wang-Claypool C.Y., Lowther, A., Parker, P., Bohec C.L. et al. (2020) Genome-wide analyses reveal drivers of penguin diversification. PNAS, 117, 22303–22310.

Waters, J.M., Fraser, C.I., Maxwell, J.J., Rawlence, N.J. (2017). Did interaction between human pressure and Little Ice Age drive biological turnover in New Zealand? Journal of Biogeography, 44, 1481–1490.

