## Supplementary Info for "Postglacial recolonization of the Southern Ocean by elephant seals occurred from multiple glacial refugia"

**SUPPORTING INFORMATION**
*Archived Archaeological and Subfossil Sites*

In order to determine the prehistoric range of elephant seals in Australasia, records of any elephant seal remains in Australia (Figure S1, Table S1) and New Zealand (Figure S2, Table S2) were collected and the minimum number of individuals documented. Locations of early Māori middens (~1250-1450 AD) containing southern elephant seal remains were sourced by accessing the archive of the New Zealand Archaeological Association Newsletter (later Archaeology in New Zealand). Records of southern elephant seal remains in Tasmanian Aboriginal middens were sourced from Bryden (1999), Stockton (1982) and Jones (1971). Additional locations of southern elephant seal remains on the Australian mainland and Tasmania were sourced from the Online Zoological Collections of Australian Museums (OZCAM, http://ozcam.org.au). Records of additional subfossil and archaeological sites containing southern elephant seals across the prehistoric and historical Southern Hemisphere range were sourced from published sources (Table S3).

*Mitochondrial Sequences from Modern Samples*

Whole genomic DNA was extracted from modern southern elephant seal tissue samples from Antipodes Island (n = 5), held in the collections of the Department of Zoology (University of Otago), using approximately 1 $mm^3$ of tissue and a 5% Chelex protocol (Walsh et al., 1991). We amplified two overlapping Control Region/D-loop segments (338 bp in total) using the primer pairs SESanc3f–SESanc3r and SESmdbf–SESmdbr (de Bruyn et al., 2009; Table S4). Each PCR reaction (10 µL) consisted of 5 µL of MyFi DNA Polymerase mix (Bioline), 0.5 µL each primer (10 mM), and 1 µL DNA. PCR thermocycling conditions consisted of 95°C 2 min, 35 cycles of 94°C 45 s, 52°C 45 s, and 72°C 45 s, with a final extension time of 72°C 10 min. PCR products were run on a 2% 1× TAE gel. Unsuccessful PCRs were repeated using 1:10 diluted DNA extract to reduce PCR inhibition. Successfully amplified PCR products were purified using ExoSAP (1.5 U ExoI, 1U SAP; GE Healthcare) with incubation at 37°C for 40 min followed by inactivation at 94°C for 15 min. PCR products were sequenced bi-directionally, using both forward and reverse primers, at Genetic Analysis Services in the Department of Anatomy, University of Otago. Contiguous Control Region/D-loop sequences were made using Sequencher 5.0 (Genecodes).

*Mitochondrial Sequences from Ancient Specimens*

We obtained 40 archaeological specimens of southern elephant seals from New Zealand and 12 from Tasmania (West Point midden, King Island), sourced from museum and university collections. New Zealand specimens were proxy dated using published radiocarbon dates on associated material and/or the faunal make-up and cultural assemblage of the site, as the large foraging range and marine reservoir effect may make the uncertainty of any radiocarbon date larger than the occupational phase at a given midden site (Petchey et al., 2008). All archaeological sites where specimens were sourced from can be dated to the 'early Māori' period of East Polynesian settlement of New Zealand (~1250-1450 AD; 750–550 BP). The Tasmanian samples from West Point midden were previously radiocarbon dated to 1-2 Kya, and the historical sample from King Island was identified as a remnant of the European sealing occupation, dating to the 19$^{th}$ Century (Ling, 1999a). Subsamples of bone and/or teeth were taken using a Dremel multitool, and a cutting disk or drill bit at low speed.

Specimens were kept in a dedicated ancient DNA (aDNA) laboratory (Otago Palaeogenetics Laboratory) physically separate from other molecular facilities (modern genetics laboratory), following strict ancient DNA procedures, including decontamination with bleach and the use of extraction, PCR, and library blanks (Knapp et al., 2012). We extracted ancient DNA following Rohland et al. (2010). We amplified up to 338 bp of Control Region/D-loop using the same primer combination as above. For samples that did not amplify for SESanc3f–SESanc3r and SESmdbf–SESmdbr, we designed new primers to amplify the 338 bp section in smaller fragments (Table S4).

Each PCR (20 µL) consisted of 5 M Betaine (Sigma), 25 mM $MgCl_2$ (Life Technologies), 10× Gold Buffer II (Life Technologies), 25 mM dNTPs (Bioline), 2 U AmpliTaq Gold DNA Polymerase (Life Technologies), 250 nM of each primer, and 2 µL of template DNA. PCR thermocycling conditions consisted of 95°C 2 min, 60 cycles of 94°C 45 s, 52°C 45 s, and 72°C 45 s, ending with 72°C 10 min. Unsuccessful PCRs were repeated with 1:10 diluted DNA to reduce PCR inhibition. Each ancient DNA extract was amplified at least twice and sequenced bi-directionally from independent PCR products to confirm sequence authenticity, alongside resolving all base ambiguities by majority-rule consensus (Brotherton et al. 2007). Contiguous Control Region/D-loop sequences were made using Sequencher 5.0 (Genecodes), producing sequences up to 325 bp long. See Table S10 for the details of the 18 successfully amplified and sequenced southern elephant seal samples.

*Mitochondrial Sequences from Ancient Bulk-Bone Assemblages*

Seersholm et al. (2018) sourced bulk samples of fragmentary, morphologically ambiguous bone (commonly termed 'frag bags' or bone-grab) from coastal early Māori archaeological midden sites around New Zealand for bulk-bone metabarcoding. DNA libraries from several archaeological sites (Fyffe Site, Tokanui, Wairau Bar, Monks Cave, Redcliffs, Saint Clair, Watsons Beach, and Awamoa) indicated the presence of southern elephant seal (Seersholm et al. 2018). Southern elephant seal mitochondrial haplotypes were determined in a two stage process: (1) the more variable of the two Control Region/D-loop fragments (233 bp) from de Bruyn et al. (2009) was amplified and sequenced from the bulk-bone libraries for the above archaeological sites, following Seersholm et al. (2018) at the Trace and Environmental DNA lab at Curtin University, Perth; (2) the same mitochondrial fragment was amplified and high-throughput sequenced from one of the successfully amplified southern elephant seal specimens following Seersholm et al. (2018) to determine the sequencing error cut-off threshold of 1.43%. Within each sequencing library, unique bulk bone haplotypes were accepted only if they were detected in at least two-thirds of replicates at abundances higher than the sequencing error cut-off threshold of >1.43% per replicate. Contiguous Control Region/D-loop sequences were made using Sequencher 5.0 (Genecodes), and combined with those from southern elephant seal specimens. Bulk-bones libraries from Fyffe Site and Watson's Beach failed to produce southern elephant seal sequence data. Up to four unique haplotypes/site were determined for Tokanui, Wairau Bar, Monks Cave, Redcliffs, Saint Clair, and Awamoa. See Tables S11-S12 for raw and summary bulk-bone metabarcoding data.

*Mitogenomes from Ancient Samples*

The 18 identifiable archaeological southern elephant seal bone samples that provided up to 325 bp of Control Region/D-loop sequences were then selected for mitochondrial genome assembly. Bulk bone data were excluded from this pool. We re-extracted ancient DNA from these selected samples in the Otago Palaeogenetics Laboratory, following Dabney et al. (2013) with 50 mg bone or tooth, which is designed to isolate ultrashort DNA fragments (>30 bp). Ancient DNA extracts (16 μL) were then prepared for hybridisation-capture enrichment of southern elephant seal DNA. The extract was first used to prepare a double-stranded DNA (dsDNA) library using the Blunt End Single Tube (BEST) method, as described by Carøe et al. (2017), using barcoded P5/P7 adapters from Gansauge and Meyer (2013). Quantitative PCR of dsDNA libraries and $C_T$ values were used to calculate the PCR number of PCR cycles for incorporation of unique Illumina adapters (Gansauge and Meyer, 2013). Each library was then amplified to incorporate a specific adapter combination in an indexing PCR. The concentration of each library was then measured with a Qubit dsDNA Broad Range assay kit (Invitrogen). For full methodology see Verry et al. (2022).

Indexed dsDNA libraries were enriched for mitochondrial DNA using baits made from a closely related species, the leopard seal (*Hydrurga leptonyx*). Genomic DNA was extracted from leopard seal tissue using the Qiagen DNeasy kit. Long-range PCR primers, specific to leopard seal, were designed to amplify the 16.5 kb mitogenome in two 8.8 kb fragments with minimal overlap. Another set of primers were also designed to amplify the mitogenome in four 4.4 kb fragments with alternative binding sites and were used when PCR amplifications with the 8.8 kb primers were unsuccessful or had low yield (Table S5). Long-range PCR reactions consisted of 2x Phusion HiFi buffer (ThermoFisher Scientific), 0.2 mM dNTPs, 0.5 μM forward and reverse long-range primers, and 0.02 U/μL HS Phusion II. We performed 50 μL reactions, with 2.5 μL of DNA extract. Thermocycling conditions were 98°C 30 s, 30 cycles of 98°C 10 s, 30 s 57–62°C gradient, and 72°C 250 s, and finally 72°C 10 min. An aliquot of each amplicon was sequenced at Macrogen Inc. (Seoul, South Korea) to confirm amplified long-range PCR products were the targeted mitochondrial regions. Amplicons were purified with the Qiagen MinElute PCR purification kit, and sheared in a Covaris E220 Evolution ultrasonicator, producing an average fragment size of 237 bp as measured on an Agilent Technologies D1000 ScreenTape. Sheared DNA concentration was measured using a Qubit dsDNA assay kit (Invitrogen), with sheared amplicons then combined in equimolar ratios. Sheared amplicons were converted to capture baits following Horn (2012), by ligating biotin oligonucleotides from Maricic et al. (2010) (Table S6). For full methodology see Verry et al. (2022).

Hybridisation capture was conducted using the biotinylated leopard seal mitogenome baits following Horn (2012), using blocking oligonucleotides designed following the same protocol. We made two changes to hybridisation conditions: a reaction volume of 80 μL, and an incubation time of 53 h. Immobilisation and purification of target-enriched libraries was performed following Horn (2012). Enriched dsDNA library concentrations were quantified using a Biorad CFX384 and the qPCR conditions from Carøe et al. (2017), in 5 μL reactions, with a 1 μL aliquot of the enriched library. IS5/6 primers from Meyer and Kircher (2010) were used in the qPCR. $C_T$ values obtained qPCR were used to calculate PCR cycle number for reamplification of enriched libraries prior to high-throughput sequencing (Gansauge and Meyer, 2013). Libraries were amplified in 25 μL triplicate reactions, using the same IS5/6 primers. Each reaction included 1.25 U AccuPrime Pfx DNA Polymerase (Invitrogen), 1× AccuPrime Pfx reaction mix, 0.3 µM primer each primer, and 7 µL of library. Libraries were then pooled, purified with a Qiagen MinElute Kit, and their concentrations measured with a Qubit dsDNA BR assay kit. Libraries were pooled in equimolar amounts and high-throughput sequenced at the Ramaciotti Centre for Genomics (University of New South Wales, Sydney) on the Illumina MiSeq 2×150bp kit.

*Analyses of Mitochondrial HVR1 Sequences*

We successfully amplified up to 338 bp of Control Region/D-loop from five modern southern elephant seals from Antipodes Island, five ancient specimens from Tasmania, and 27 ancient individuals (from a combination of specimens and bulk bone assemblages; 18 individual specimens and nine unique haplotypes from bulk bone libraries from six different archaeological sites) from New Zealand. We combined these with 591 publicly available Control Region/D-loop sequences to produce a data set containing representatives from all known populations of southern elephant seal (Table 1) (Hoelzel et al., 1993; Slade et al., 1998; Fabiani et al., 2003; de Bruyn et al., 2009; Bogdanowicz et al., 2013; Corrigan et al., 2016).

All statistical analyses were carried out with controls for heterochronous data recommended by Depaulis et al. (2009). We used DnaSP v6 (Rozas et al., 2017) to evaluate genetic diversity at each location, using a number of metrics: number of haplotypes (*n*); number of segregating sites (*S*); average nucleotide differences per sequence (*k*); haplotype diversity (*h/n*); and nucleotide diversity with the Jukes-Cantor correction ($\pi JC$). From this data, we also calculated an 'expansion coefficient' (*S/k*) to test for an increase in population size over time (Peck and Congdon, 2004; de Bruyn et al., 2009) (Table 1).

We calculated $F_{ST}$ values using Arlequin v3.5 (Excoffier and Lischer, 2010) to quantify levels of genetic differentiation between population pairs. Fixation values, geographic data, and genetic clades found in previous studies were then used to construct groups of breeding stocks. We performed analyses of molecular variance (AMOVAs) in Arlequin.

The dataset was trimmed to the 233 bp of Control Region common to modern and archaeological (identifiable specimens and bulk bone metabarcoding libraries). Specimens with too little sequence data (see Table X) were removed from the analysis (three from New Zealand and two from Tasmania). We then phylogenetically analysed the Control Region/D-loop sequences to construct a time-sensitive haplotype network in TempNet (Prost et al., 2011) and PopArt (Figure S4). Identical haplotypes that occur at different time points are linked, showing relatedness of ancient and modern samples (Figure 3).

*Analyses of Whole Mitochondrial Genomes*

High-throughput sequence data was processed using the "trim_merge_DS_PE_standard.sh" and "map_SE.sh" scripts within the BEARCAVE data analysis and storage environment (https://github.com/nikolasbasler/BEARCAVE), which are publicly available and can be used to replicate the analysis. This pipeline involved trimming adapter sequences from read pairs using the program Cutadapt v1.12 (Martin, 2011) and removing any pairs < 30 bp after trimming, using a minimum adapter sequence overlap of 1 bp and default values for all other parameters. Overlapping read pairs were then identified and merged using the program FLASH v1.2.11 (Magoč & Salzberg, 2011) using default parameter values. Only successfully merged read pairs, which are effectively single-end data, were used for mapping.

Reads were mapped to the southern elephant seal reference mitochondrial genome (GenBank accession no.: NC_008422), using the aln algorithm of the program bwa v0.7.15 (Li & Durbin, 2009) with default parameter values, and then filtered for reads with low mapping quality (-q 30) and sorted by 5' mapping position using the program samtools v1.3.1 (Li et al., 2009). Potential PCR duplicates were then identified from resulting bam files using the program MarkReadsByStartEnd.jar (https://github.com/dariober/Java-cafe/tree/master/MarkDupsByStartEnd), and removed using samtools. Consensus sequences were then computed in Geneious v7.0, applying a minimum depth of 3 reads and a 75% majority rule for base calling. The program MapDamage v2.0.8 (Ginolhac et al., 2011) was used to assess whether the data exhibits patterns of damage typical for ancient DNA.

The results of data processing are shown in Table S7 and consensus calling in Table S8. Overall, we were able to recover partial mitochondrial genome sequences for 10 of the 18 specimens (with an average depth cutoff of 2X), with a mean depth of coverage of 2.28–22.35 reads, and a percentage recovery of the total mitochondrial genome of 46.8–96.4 %. The MapDamage analyses were affected by the generally low numbers of mapped reads. Although the data were generally noisy, indicative patterns of C→T and G→A substitutions could be evidenced for all specimens at the 5' and 3' fragment ends, respectively, indicating authentic ancient DNA. Although the accurate recovery of DNA fragment length distributions was hampered by the low number of sampled reads, DNA fragmentation typical of ancient DNA was also evident for all specimens.

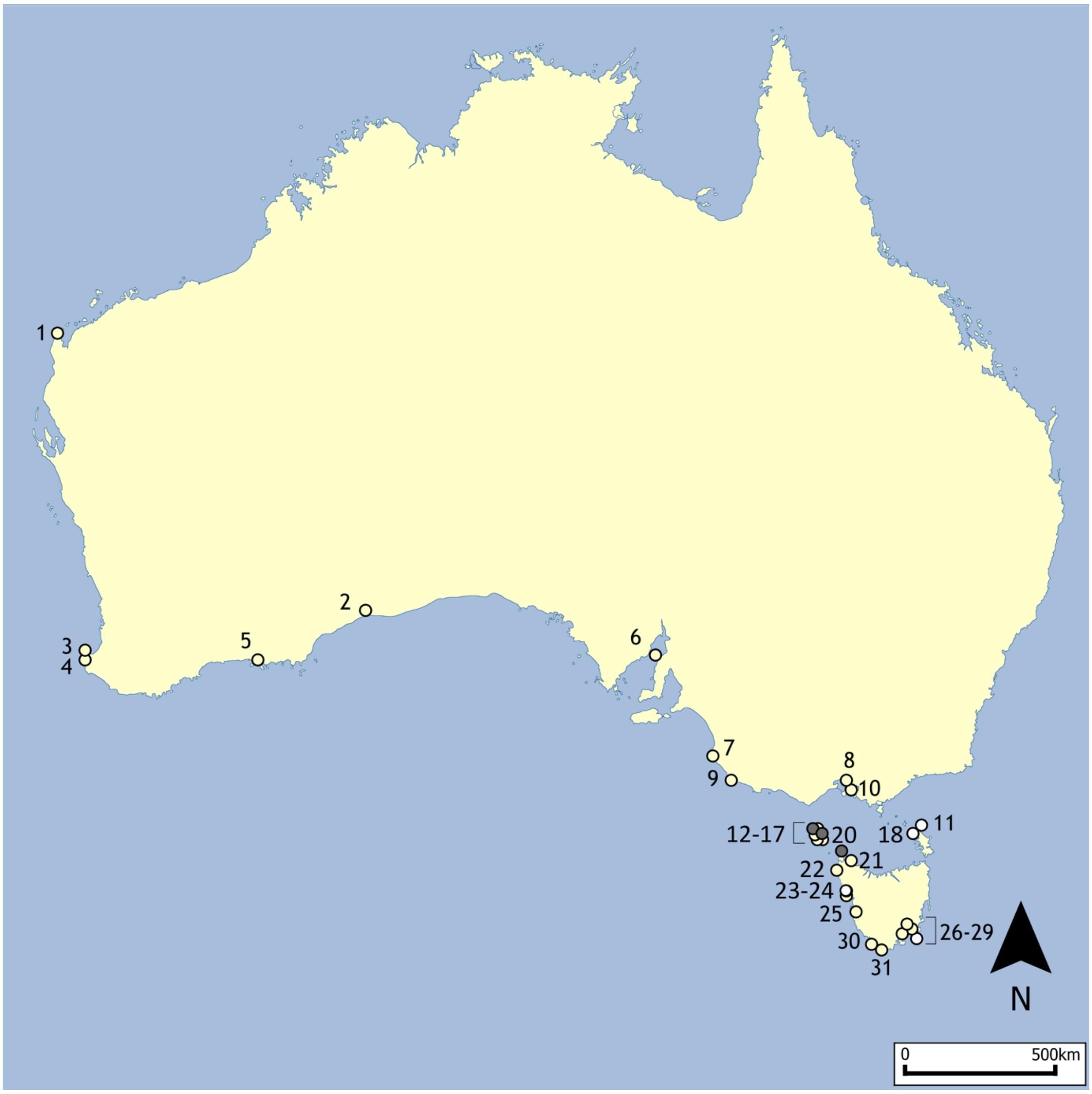

**Figure S1:** Australian archaeological records of southern elephant seals. Location details (numbered sites), minimum number of individuals and site ages are in Table S1. White circles indicate the presence of southern elephant seal remains; dark grey circles indicate historically exploited populations.

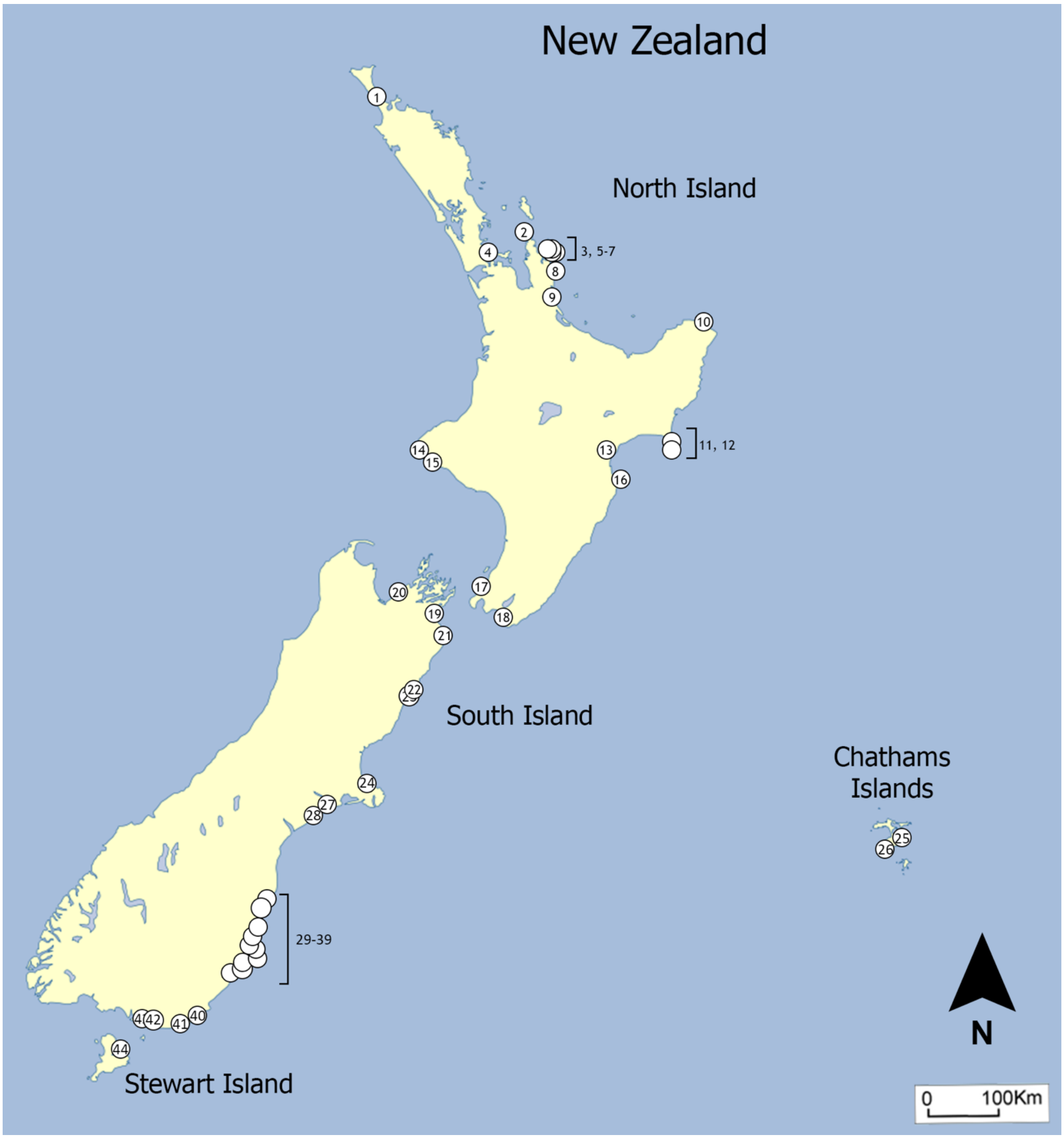

**Figure S2:** Aotearoa New Zealand early Maori (1250-1450 AD) archaeological records of identifiable southern elephant seals (not including bulk-bone remains). Location details (numbered sites) and minimum number of individuals are in Table S2.

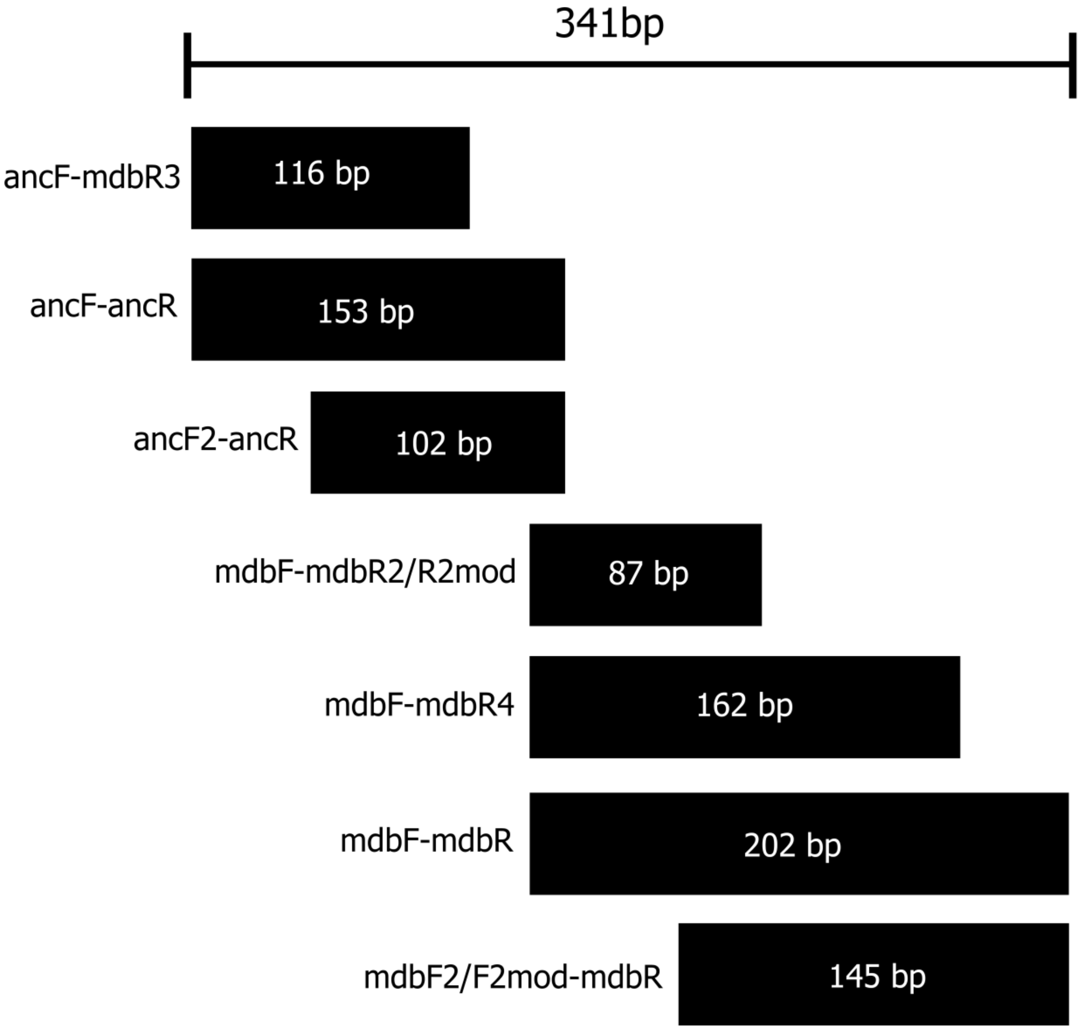

**Figure S3:** The total length of the southern elephant seal mitochondrial DNA (mtDNA) Control Region/D-loop fragment used in this study (excluding primers), originally from de Bruyn et al. (2009), including primer pairs and the length of amplicon (excluding primers).

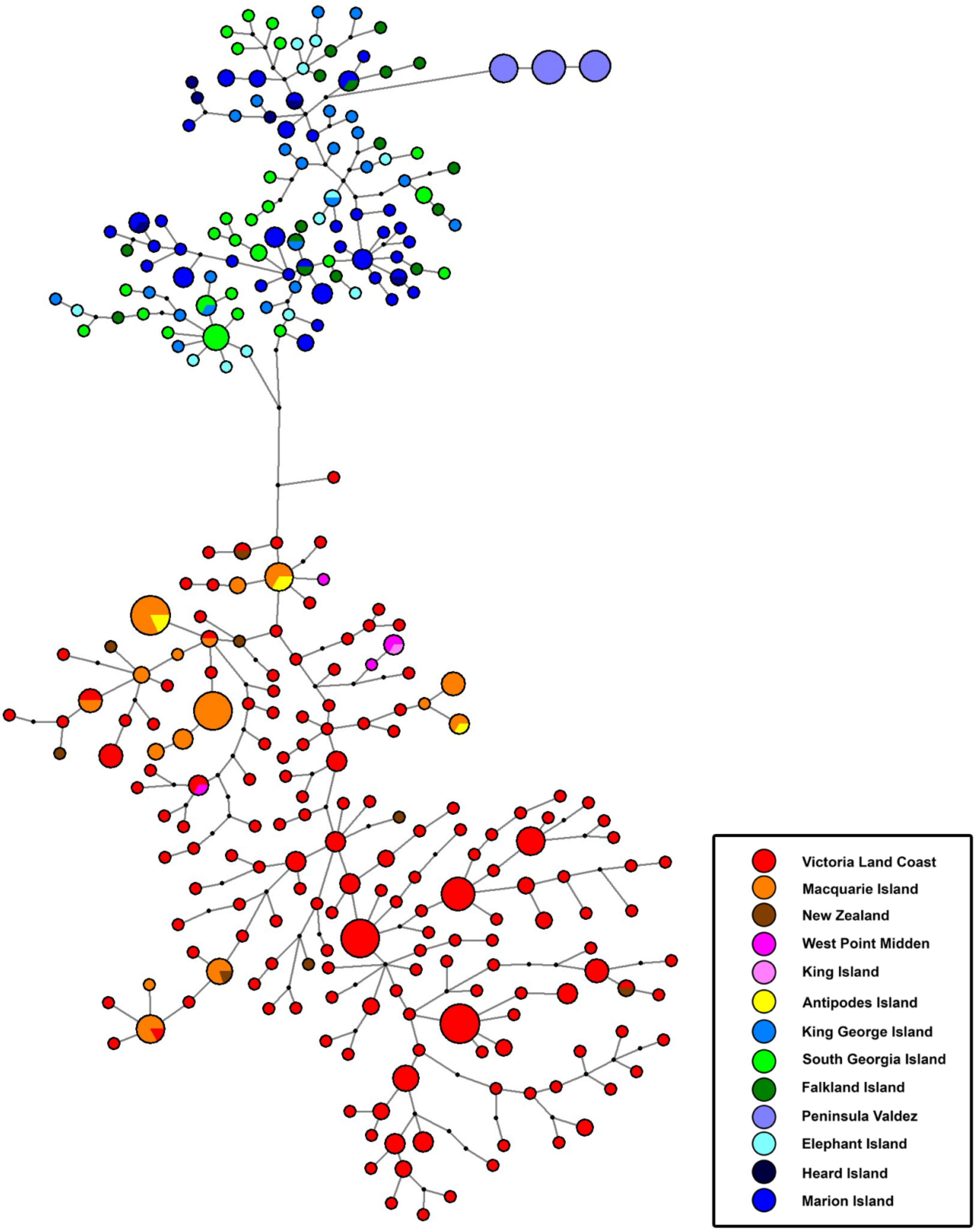

**Figure S4:** A median-joining network using 233 bp of the Control Region/D-loop of southern elephant seal representing the whole breeding range of the species, with all ancient samples included but inferred migrants removed (Table S9). Haplogroup size is indicative of frequency; branch lengths do not reflect sequence differences. Colours represent sampling locations (inset). Small black circles represent median vectors.

**Table S1:** The location of southern elephant seals on the Australian mainland, Bass Strait (and its associated islands) and Tasmania; each separated by double lines. Abbreviations: minimum number of individuals (MNI) where known, otherwise indicative numbers based on literature. The map numbers correspond to those in Figure S1.

| Map #: | Location: | MNI: | Sample age: | Reference: |
|---|---|---|---|---|
| 1 | North West Cape Area, Western Australia | 1 | Unknown | OZCAM |
| 2 | Dundas, Western Australia | 1 | Unknown | OZCAM |
| 3 | Yalingup Beach, Western Australia | 1 | Unknown | OZCAM |
| 4 | Moses Rock, Western Australia | 1 | Unknown | OZCAM |
| 5 | Wylie Bay, Western Australia | 1 | Unknown | OZCAM |
| 6 | Spencer Gulf, South Australia | 1 | Unknown | OZCAM |
| 7 | Robe, South Australia | 1 | Unknown | OZCAM |
| 8 | Parkdale Beach, Victoria | 1 | Unknown | OZCAM |
| 9 | Port Macdonnell, South Australia | 1 | Unknown | OZCAM |
| 10 | Seal Rocks, Victoria | 1 | Unknown | OZCAM; Warneke (1976) |
| 11 | East Sister Island | Rare individuals | 19th Century | Warneke (1976); Stockton (1982) |
| 12-16 | King Island (incl. Sea Elephant Bay) | Large numbers | 19th Century | OZCAM; Micco (1971); Warneke (1976); Stockton (1982) |
| 17 | Cuvier Point, King Island | 1 | Mid-Late Holocene | Stockton (1982) |
| 18 | New Year Island | Large numbers | 19th Century | Micco (1971); Warneke (1976) |
| 19 | North East Point, Flinders Island | 1 | 19th Century | Warneke (1976) |
| 20 | Hunter/Barren Islands | Large numbers | 19th Century | Micco (1971); Warneke (1976) |
| 21 | Lee Archer, Rocky Cape, Sisters Creek, Tasmania | 11 | 8000ybp-400ybp | Jones (1971) |
| 22 | West Point, Tasmania | >1000 | 2000-1000 ybp | Bryden et al. (1999) |
| 23 | Sundown Point, Tasmania | mentioned | Mid-Late Holocene | Stockton (1982) |
| 24 | Nelson Bay, Tasmania | mentioned | Mid-Late Holocene | Stockton (1982) |
| 25 | Sloop Rocks, Tasmania | mentioned | Mid-Late Holocene | Stockton (1982) |
| 26 | Maria Point, Tasmania | 1 | Unknown | OZCAM |
| 27 | Sandy Bay, Tasmania | 1 | Unknown | OZCAM |
| 28 | Mable Bay, Tasmania | 1 | Unknown | OZCAM |
| 29 | Lighthouse Bay, Tasmania | 1 | Unknown | OZCAM |
| 30 | Maatsuyuker Is., Tasmania | mentioned | Mid-Late Holocene | Stockton (1982) |
| 31 | Louisa Bay, Tasmania | mentioned | Mid-Late Holocene | Stockton (1982) |

**Table S2:** The location of early Maori (1250-1450 AD) archaeological midden and subfossil (in bold) deposits containing southern elephant seal remains in the New Zealand region. Archaeological sites have been dated to this period using a combination of radiocarbon dating associated material and/or cultural assemblage. Abbreviations: minimum number of individuals (MNI). Double lines separate the main islands of New Zealand. The map numbers listed correspond to those in Figure S2.

| Map #: | Location: | MNI: | Reference: |
|---|---|---|---|
| 1 | Houhora, North Island | 8 | Smith (1985) |
| 2 | Port Jackson, North Island | 1 | Smith (2013) |
| 3 | Cross Creek, North Island | 1 | Smith (2013) |
| 4 | Sunde, North Island | only mentioned | Smith (2013) |
| 5 | Sarahs Gully, North Island | 1 | Smith (1985) |
| 6 | Opito Beach, North Island | 2 | Smith (1985) |
| 7 | Skippers Ridge, North Island | only mentioned | Smith (1985) |
| 8 | Tairua, North Island | 1 | Smith (2013) |
| 9 | Whangamata, North Island | only mentioned | Smith (1985) |
| 10 | Onepoto Beach | only mentioned | Smith (1985) |
| 11 | Onenui, North Island | only mentioned | Smith (1985) |
| 12 | Mahia Peninsula, North Island | 1 | Jeal (1987); Jones et al. (2003) |
| 13 | Waipunga, North Island | 1 | Boessenecker and Churchill (2016) |
| 14 | Opua, North Island | 1 | Fyfe (1988) |
| 15 | Kaupokonui, North Island | 1 | Smith (1985) |
| 16 | Rangaiika, North Island | only mentioned | Smith (1985) |
| 17 | Paremata, North Island | 4 | Smith (1985) |
| 18 | Washpool, North Island | 2 | Smith (1985) |
| 19 | Wairau Bar, South Island | only mentioned | Smith (1985) |
| 20 | Rotokura, South Island | 5 | Smith (1985) |
| 21 | Avoca Point, South Island | only mentioned | Smith (1985) |
| 22 | Marfells Beach, South Island | only mentioned | Smith (1985) |
| 23 | Fyffe Site, South Island | | Seersholm et al. (2018) |
| 24 | Redcliffs, South Island | 3 | Challis (1995) |
| 25 | Hanson Bay, Chatham Islands | only mentioned | Smith and Wernham (1976) |
| 26 | Point Durham, Chatham Islands | 12 | Sutton and Marshall (1980) |
| 27 | Rakaia Mouth, South Island | only mentioned | Smith (1985) |
| 28 | Wakanui, South Island | 1 | Challis (1995) |
| 29 | Waitaki River Mouth, South Island | 1 | Challis (1995) |
| 30 | Waimataitai Mouth, South Island | 1 | Smith (2013) |
| 31 | Awamoa, South Island | | Seersholm et al. (2018) |
| 32 | Shag River Mouth, South Island | 17-34 | Nagaoka (2000) |
| 33 | Pleasant River Mouth, South Island | 6-8 | Smith (1985, 1999) |
| 34 | Purakanui, South Island | 1 | Smith (1985) |
| 35 | Long Beach, South Island | 2 | Smith (2013) |
| 36 | Hoopers Inlet, South Island | only mentioned | Smith (1985) |
| 37 | St. Clair, South Island | 2 | Jacomb et al. (2010) |
| 38 | Otokia Mouth, South Island | 1 | Anderson (1982) |
| 39 | Watsons Beach, South Island | | |
| 40 | Pounawea, South Island | 2 | Smith (2013) |
| 41 | Papatowai, South Island | 2 | Smith (2013) |
| 42 | Tokanui River Mouth, South Island | | Seersholm et al. (2018) |
| 43 | Tiwai Point, South Island | 6 | Sutton and Marshall (1980) |

| | | | |
|---|---|---|---|
| 44 | Ringaringa, Rakiura Stewart Island | 2 | Knight (1970) |

**Table S3:** The location of archaeological and subfossil southern elephant seal remains in the southern hemisphere. Abbreviations: minimum number of individuals (MNI).

| Location: | MNI: | Sample age: | Reference: |
|---|---|---|---|
| Norfolk Island | 1 | 0.8-0.6 Kya | Anderson (2001); Anderson and White (2001) |
| Raoul Island | 1 | 0.7-0.6 Kya | Anderson (2001); Anderson and White (2001) |
| Juan Fernandez | present | 200-100 ybp | Ling and Bryden (1992) |
| Antofagasta, South America | 1 | 330 kya | Valenzuela-Toro et al., (2015) |
| Kasteelberg, South Africa | 1 | 1.6 kya | Smith (2006); Avery and Klein (2011) |
| Nelson Bay Cave, South Africa | 2 | 10-8 kya | Klein (1972a, b); Avery and Klein (2011) |
| Ysterfontein, South Africa | 1 | <2 kya | Hendey (1974); Avery and Klein (2011); |
| Die Kelders Cave 1, South Africa | 1 | 1.6 kya | Schweitzer (1979); Klein and Cruz-Uribe (2000); Avery and Klein (2011) |
| Victoria Land Coast, Antarctica | >1000 | 8-1 kya | De Bruyn et al. (2009); Hall et al. (2023) |

**Table S4**: Primers used to amplify ancient and modern Control Region/D-loop fragments of southern elephant seal.

| Primer name | Sequence | Reference |
|---|---|---|
| ancF | 5’-GCTGACATTCTACTTAAACT-3’ | de Bruyn et al., (2009) |
| ancR | 5’-ATGTACATGCTTATATGCAT-3’ | de Bruyn et al., (2009) |
| mdbF | 5’-AGCCCTATGTATATCCTGCATT-3’ | de Bruyn et al., (2009) |
| mdbR | 5’-CAGTATAGAAACCCCCACATGA-3’ | de Bruyn et al., (2009) |
| mdbF2 | 5’-ACTGGTRTGATTTTACATAAT-3’ | This study |
| mdbF2mod | 5’-ACTGGTTGATTTTACATRAT-3’ | This study |
| mdbR2 | 5’-GCATYCATTATAGGTAGTTTTA-3’ | This study |
| mdbR2mod | 5’-GCATYCATTATAGGTAGTTTTA-3’ | This study |
| mdbR3 | 5’-AATGCAGGATATACATAGGGCT-3’ | This study |
| mdbR4 | 5’-GATTACGTACACGTTTCACAA-3’ | This study |
| ancF2 | 5’-ATATAACATCACTTYCACTGT-3’ | This study |

**Table S5**: Long-range PCR primers designed to amplify leopard seal mitochondrial genomes.

| Primer name | Sequence |
|---|---|
| 9kb-1F | 5’-ACAGTATGAACRATCCTACCAGC-3’ |
| 9kb-1R | 5’-ACTGCGWCGAGACCTTTACG’-3’ |
| 9kb-2F | 5’-YCAGCAACCCTTGTGAAACG-3’ |
| 9kb-2R | 5’-TGGGATGGCRTCGGTTTTTA-3’ |
| 4kb-1F | 5’-ACCAGTTCCTCCCAGTACGA-3’ |
| 4kb-1R | 5’-CGTGTGGTCGTGGAAGTRTA-3’ |
| 4kb-2F | 5’-CCTCYATGGCGTACCCTCTCC-3’ |
| 4kb-2R | 5’-GTGACRAAGAGGGCTACRGG-3’ |
| 4kb-3F | 5’-AGCTCAAACCTTTTATTTACCGA-3’ |
| 4kb-3R | 5’-ACTGCGWCGAGACCTTTACG-3’ |
| 4kb-4F | 5’-CCAGCAACCCTTGTGAAACG-3’ |
| 4kb-4R | 5’-GGTCCTACGATGTTGGGTCC-3’ |

**Table S6**: Blocking and biotin oligonucleotides synthesised for whole mitogenome hybridisation capture.

| Oligonucleotide name | Sequence | Reference |
| --- | --- | --- |
| Bio-T | 5’-Biotin-TCAAGGACATCCG-3’ | Maricic et al., 2010 |
| B | 5’-CGGATGTCCTTG-3’ | Maricic et al., 2010 |
| Block_P7 | 5’-CAAGCAGAAGACGGCATACGAGATNNNNNNNGTGACTGGAGTTCA GACGTGTGCTCTTCCGATCT-Phosphate-3’ | This paper (adapted from Horn, 2012) |
| Block_P5 | 5’-AATGATACGGCGACCACCGAGATCTACACNNNNNNNACACTCTTTC CCTACACGACGCTCTTCCGATCT-Phosphate-3’ | This paper (adapted from Horn, 2012) |

**Table S7**: Results of mitochondrial genome data processing. Abbreviations are as follows: KI: King Island, Tasmania; WPM: West Point Midden, Tasmania; HNZ: Houhora, New Zealand; PAMZ: Paremata, New Zealand; WBNZ: Wairau Bar, New Zealand; RCNZ: Redcliffs, New Zealand; TBNZ: Tumbledown Bay, New Zealand; SMNZ: Shag River Mouth, New Zealand; PONZ: Pounawea, New Zealand; NTC23 and NTC28: No Template Controls.

| Specimen | Location | n_reads | n_under30bp | prop_under30bp | n_over30bp | n_merged | prop_merge | n_readsformapping | n_mapped | n_uniq | prop_duplicates | depth | n_mappedbp |
|---|---|---|---|---|---|---|---|---|---|---|---|---|---|
| NRO45 | KI | 894817 | 59601 | 0.07 | 835216 | 727723 | 0.87 | 727723 | 882 | 407 | 0.54 | 4.35 | 68099 |
| Lfemale03-085 | WPM | 1066830 | 57267 | 0.05 | 1009563 | 929883 | 0.92 | 929883 | 281 | 261 | 0.07 | 2.28 | 29556 |
| LfemaleF3 | WPM | 875352 | 302201 | 0.35 | 573151 | 528926 | 0.92 | 528926 | 369 | 122 | 0.67 | 1.51 | 13127 |
| LfemaleN2 | WPM | 1159004 | 30515 | 0.03 | 1128489 | 1068386 | 0.95 | 1068386 | 1212 | 716 | 0.41 | 5.93 | 96020 |
| RfemaleK1B | WPM | 669477 | 21476 | 0.03 | 648001 | 596736 | 0.92 | 596736 | 382 | 36 | 0.91 | 1.15 | 3735 |
| LmaleL5a | WPM | 144464 | 14771 | 0.10 | 129693 | 117992 | 0.91 | 117992 | 625 | 187 | 0.70 | 1.56 | 12882 |
| RmaleL5 | WPM | 953627 | 51723 | 0.05 | 901904 | 834772 | 0.93 | 834772 | 62 | 59 | 0.05 | 1.28 | 8357 |
| RmaleL5a3 | WPM | 1468066 | 63222 | 0.04 | 1404844 | 1348501 | 0.96 | 1348501 | 487 | 422 | 0.13 | 2.76 | 40959 |
| NRO287 | HNZ | 800541 | 53804 | 0.07 | 746737 | 725762 | 0.97 | 725762 | 24 | 23 | 0.04 | 1.08 | 2655 |
| NRO290 | HNZ | 927068 | 45976 | 0.05 | 881092 | 829883 | 0.94 | 829883 | 177 | 155 | 0.12 | 1.66 | 16485 |
| NRO156 | PANZ | 897875 | 23035 | 0.03 | 874840 | 839792 | 0.96 | 839792 | 11 | 7 | 0.36 | 1.05 | 591 |
| NRO66 | WBNZ | 1110183 | 91971 | 0.08 | 1018212 | 997922 | 0.98 | 997922 | 92 | 89 | 0.03 | 1.30 | 7859 |
| WB01 | WBNZ | 3276131 | 179223 | 0.05 | 3096908 | 2905306 | 0.94 | 2905306 | 1366 | 1239 | 0.09 | 8.86 | 146044 |
| NRO39 | RCNZ | 1105691 | 14804 | 0.01 | 1090887 | 1048824 | 0.96 | 1048824 | 8 | 8 | 0.00 | 1.07 | 741 |
| TB16H3 | TBNZ | 2292393 | 130547 | 0.06 | 2161846 | 1922878 | 0.89 | 1922878 | 21312 | 2778 | 0.87 | 22.35 | 368217 |
| BQ546 | SMNZ | 975955 | 77587 | 0.08 | 898368 | 866622 | 0.96 | 866622 | 25926 | 2815 | 0.89 | 19.07 | 314191 |
| SMC-BQ-092-1 | SMNZ | 1281574 | 175435 | 0.14 | 1106139 | 1041229 | 0.94 | 1041229 | 7 | 6 | 0.14 | 1.02 | 599 |
| NRO38 | PONZ | 1020214 | 51964 | 0.05 | 968250 | 904662 | 0.93 | 904662 | 104 | 81 | 0.22 | 1.29 | 8092 |
| NTC23 | - | 1207941 | 919735 | 0.76 | 288206 | 8389 | 0.03 | 8389 | 12 | 12 | 0.00 | 1.05 | 1439 |
| NTC28 | - | 821601 | 592839 | 0.72 | 228762 | 7057 | 0.03 | 7057 | 11 | 11 | 0.00 | 1.04 | 1494 |

**Table S8:** GenBank accession numbers of putative migrant individuals, showing recorded location, likely location of origin, and the basis of inferred migrant status.

| GenBank accession number | Location in which the individual was recorded | Likely location of origin | Migrant identification method | Reference |
|---|---|---|---|---|
| JX847057 | King George Island | Unknown | GENECLASS2 analysis | Bogdanowicz et al., (2013) |
| JX847044 | King George Island | Unknown | GENECLASS2 analysis | Bogdanowicz et al., (2013) |
| AY166627 | Falkland Island | Macquarie Island | Shared mtDNA haplotype | Fabiani et al., (2003) |
| AY769723 | Marion Island | Macquarie breeding area | Shared mtDNA haplotype | Chauke (2008) |
| DQ267956 | Marion Island | Macquarie breeding area | Shared mtDNA haplotype | Chauke (2008) |

**Table S9**: Mitochondrial Control Region $F_{ST}$ values for southern elephant seal populations using the Kimura 2P distance method, 10,000 permutations. Bolded values have significant P-values (*p*<0.05). Grey shading indicates comparisons between Australasian sequences. NZ = New Zealand, TAS = Tasmania, VLC = Victoria Land Coast, ANT = Antipodes Islands, MQ = Macquarie Island, KGI = King George Island, MI = Marion Island, SG = South Georgia, HI = Heard Island, EI = Elephant Island, FI = Falkland Islands, PV = Peninsula Valdés.

| | NZ | TAS | VLC | ANT | MQ | KGI | MI | SG | HI | EI | FI |
|---|---|---|---|---|---|---|---|---|---|---|---|
| **TAS** | **0.24204** | - | - | - | - | - | - | - | - | - | - |
| **VLC** | **0.19064** | **0.4297** | - | - | - | - | - | - | - | - | - |
| **ANT** | 0.0442 | **0.5109** | **0.2208** | - | - | - | - | - | - | - | - |
| **MQ** | 0.04472 | **0.2591** | **0.2164** | -0.0303 | - | - | - | - | - | - | - |
| **KGI** | **0.23957** | 0.1042 | **0.4628** | **0.37198** | **0.3029** | - | - | - | - | - | - |
| **MI** | **0.33228** | **0.198** | **0.5111** | **0.43842** | **0.3759** | **0.0639** | - | - | - | - | - |
| **SG** | **0.32678** | **0.2385** | **0.5164** | **0.43324** | **0.3622** | **0.09097** | **0.16226** | - | - | - | - |
| **HI** | **0.28927** | **0.2958** | **0.4895** | **0.48449** | **0.3021** | 0.0026 | 0.0291 | **0.17896** | - | - | - |
| **EI** | **0.28555** | 0.1144 | **0.4831** | **0.39932** | **0.3288** | 0.0046 | **0.11738** | **0.08907** | 0.08274 | - | - |
| **FI** | **0.31107** | **0.1916** | **0.486** | **0.40816** | **0.3696** | **0.03241** | **0.04491** | **0.15905** | -0.0325 | **0.08374** | - |
| **PV** | **0.73926** | **0.924** | **0.6553** | **0.9448** | **0.6334** | **0.56796** | **0.4989** | **0.56698** | **0.74739** | **0.62581** | **0.36503** |

**Table S10:** Southern elephant seal specimens for which mitochondrial sequence data was generated in this study. * Bulk bone metabarcoding by Seersholm et al. (2018) indicated the presence of southern elephant seal 16S rRNA but failed to generate unique CR haplotypes in this study. # produced mtDNA CR but not mitogenome sequence. Grey shaded specimens produced fragmentary CR data so were excluded from the population genetic and haplotype network analysis.

| Specimen name | Geographic Location | Museum/University | Colln # |
|---|---|---|---|
| *Modern specimens: mtDNA CR* | | | |
| SES1 | Antipodes Island | University of Otago (Zoology) | SES1 |
| SES2 | Antipodes Island | University of Otago (Zoology) | SES2 |
| SES3 | Antipodes Island | University of Otago (Zoology) | SES3 |
| SES4 | Antipodes Island | University of Otago (Zoology) | SES4 |
| SES6 | Antipodes Island | University of Otago (Zoology) | SES6 |
| *Bulk bone metabarcoding aDNA libraries from Early Maori archaeological sites in NZ (Seersholm et al. 2018): mtDNA CR* | | | |
| Awamoa | Awamoa | University of Otago (Zoology) | |
| Monks Cave | Redcliffs | Canterbury Museum | |
| Main Road 27 | Redcliffs | University of Otago (Archaeology) | |
| Saint Clair | Saint Clair | University of Otago (Archaeology) | |
| Tokanui | Tokanui | University of Otago (Archaeology) | |
| Wairau Bar | Wairau Bar | University of Otago (Archaeology), Canterbury Museum | |
| Fyffe Site* | Kaikoura | Canterbury Museum | |
| Watsons Beach* | Watsons Beach | University of Otago (Archaeology) | |
| *Identifiable southern elephant seals from Early Maori archaeological sites in NZ: mtDNA CR and mitogenome* | | | |
| NRO287 | Houhora | Auckland Museum | AM unregistered |
| NRO290 | Houhora | Auckland Museum | AM unregistered |
| NRO156 | Paremata | Museum of New Zealand Te Papa Tongawera | BG 44/2 |
| WB01 | Wairau Bar | Canterbury Museum | CM unregistered |
| NRO66 | Wairau Bar | Canterbury Museum | CM Ma1325 |
| NRO39 | Redcliffs | Canterbury Museum | CM Ma1145 |

| | | | |
|---|---|---|---|
| TB16H3 | Tumbledown Bay | Brian Allingham collection | TB16H3 |
| TB17G2[#] | Tumbledown Bay | Brian Allingham collection | TB17G2 |
| BQ692 | Shag River Mouth | University of Otago (Archaeology) | BQ692 |
| BQ546_10 | Shag River Mouth | University of Otago (Archaeology) | BQ546_10 |
| NRO38 | Pounawea | Canterbury Museum | CM Ma3424 |
| *Identifiable archaeological southern elephant seals from Tasmania: mtDNA CR and mitogenome* | | | |
| NRO45 | King Island | Canterbury Museum | CM Ma640 |
| L♂L5a | West Point | Australian National University | L♂L5a |
| R♂L5 | West Point | Australian National University | R♂L5 |
| R♂L5a3 | West Point | Australian National University | R♂L5a3 |
| L♀N2 | West Point | Australian National University | L♀N2 |
| L♀F3 | West Point | Australian National University | L♀F3 |
| L♀03 | West Point | Australian National University | L♀03 |
| R♀K1B 053 | West Point | Australian National University | R♀K1B 053 |

**Supplementary References**

Anderson, A. (1982) The Otokia Mouth site at Brighton Beach, Otago. *New Zealand Archaeological Association Newsletter,* 47-52.

Anderson, A. (2001) No meat on that beautiful shore: the prehistoric abandonment of subtropical Polynesian islands. *International Journal of Osteoarchaeology*, 11, 14-23.

Anderson, A., White, P. (Eds) (2001). The prehistoric archaeology of Norfolk Island, Southwest Pacific. Records of the Australian Museum, Supplement 27.

Avery, G., Klein, R.G. (2011) Review of fossil phocid and otariid seals from the southern and western coasts of South Africa. *Transactions of the Royal Society of South Africa*, 66, 14-24.

Bogdanowicz, W., Pilot, M., Gajewska, M., Suchecka, E., Golachowski, M. (2013) Genetic diversity in a moulting colony of southern elephant seals in comparison with breeding colonies. *Marine Ecology Progress Series*, 478, 287–300.

Bryden, M.M., O'Connor, S., Jones, R. (1999) Archaeological evidence for the extinction of a breeding population of elephant seals in Tasmania in prehistoric times. *International Journal of Osteoarchaeology,* 9**,** 430–437.

Carøe, C., Gopalakrishnan, S., Vinner, L., Mak, S.T.M., Sindig, M.H.S., Samaniego, J.A., Wales, N., Sicheritz-Pontén, T., Gilbert, M.T.P. (2018) Single-tube library preparation for degraded DNA. *Methods in Ecology and Evolution*, 9, 410–419.

Challis, A.J. (1995) *Ka pakihi whakatekateka o waitaha: the archaeology of Canterbury in Maori times*: Department of Conservation Wellington.

Corrigan, L.J., Fabiani, A., Chauke, L., McMahon, C.R., de Bruyn, M., Bester, M.N., Bastos, A., Campagna, C., Muelbert, M.M., Hoelzel, A.R. (2016) Population differentiation in the context of Holocene climate change for a migratory marine species, the southern elephant seal. *Journal of Evolutionary Biology,* 29, 1667–1679.

Dabney, J., Knapp, M., Glocke, I., Gansauge, M. T., Weihmann, A., Nickel, B., Valdiosera, C., Garcia, N., Paabo, S., Arsuaga, J. L. Meyer, M. (2013) Complete mitochondrial genome sequence of a Middle Pleistocene cave bear reconstructed from ultrashort DNA fragments. *Proceedings of the National Academy of Sciences of the United States of America,* 110**,** 15758–15763.

de Bruyn, M., Hall, B.L., Chauke, L.F., Baroni, C., Koch, P.L., Hoelzel, A.R. (2009). Rapid response of a marine mammal species to Holocene climate and habitat change. *PLOS Genetics, 5.*

Depaulis, F., Orlando, L., Hänni, C. (2009) Using classical population genetics tools with heterochroneous data: time matters!. *PLOS ONE*, 4, e5541.

Excoffier, L., Lischer, H.E. (2010) Arlequin suite ver 3.5: a new series of programs to perform population genetics analyses under Linux and Windows. *Molecular Ecology Resources,* 10**,** 564–567.

Fabiani, A., Hoelzel, A. R., Galimberti, F., Muelbert, M.M.C. (2003) Long-range paternal gene flow in the southern elephant seal. *Science,* 299**,** 676–676.

Gansauge, M.T., Meyer, M. (2013) Single-stranded DNA library preparation for the sequencing of ancient or damaged DNA. *Nature Protocols,* 8**,** 737–748.

Grealy, A.C., McDowell, M.C., Scofield, P., Murray, D.C., Fusco, D.A., Haile, J., Prideaux, G.J., Bunce, M. (2015) A critical evaluation of how ancient DNA bulk bone metabarcoding complements traditional morphological analysis of fossil assemblages. *Quaternary Science Reviews*, 128, 37–47.

Hall, B.L., Koch, P.L., Baroni, C., Salvatore, M.C., Hoelzel, A.R., de Bruyn, M., Welch, A. (2023) Widespread southern elephant seal occupation of the Victoria Land Coast implies a warmer-than-present Ross Sea in the mid-to-late Holocene. *Quaternary Science Reviews*, 303, 107991.

Hendey, Q.B. (1974) Faunal dating of the late Cenozoic of southern Africa, with special reference to the Carnivora. *Quaternary Research*, 4, 149-161.

Hoelzel, A., Halley, J., O'Brien, S.J., Campagna, C., Arnborm, T., Le Boeuf, B., Ralls, K., Dover, G. (1993) Elephant seal genetic variation and the use of simulation models to investigate historical population bottlenecks. *Journal of Heredity,* 84**,** 443–449.

Horn, S. (2012) Target enrichment via DNA hybridization capture. *In:* Shapiro, B., Hofreiter, M. (eds) *Ancient DNA: Methods and Protocols.* New York: Springer.

Jacomb, C., Brooks, E., Mintmier, M., Walter, R. (2010) Final report on archaeological investigations at I44/121, St Blair, Dunedin. *Unpublished report for St Clair Project Limited by South Pacific Archaeological Research*.

Jeal, M. (1987) A Mahia Peninsula sea mammal butchery site (N127/14). *New Zealand Archaeological Association Newsletter*, 174-175.

Jones, K.L., Jeal, M., Jeal, M. (2003) Field archaeology of the Mahia Peninsula (Nukutaurua mai Tawhiti), northern Hawkes Bay, New Zealand. *New Zealand Journal of Archaeology*, 23, 5-29.

Jones, R. (1971) *Rocky Cape and the problem of the Tasmanians*. University of Sydney. PhD Thesis.

Klein, R.G. (1972a) The late Quaternary mammalian fauna of Nelson Bay Cave (Cape Province, South Africa): its implications for megafaunal extinctions and environmental and cultural change. *Quaternary Research*, 2, 135-142.

Klein, R.G. (1972b). Preliminary report on the July through September excavations at Nelson Bay Cave, Plettenberg Bay (Cape Province, South Africa). *Palaeoecology of Africa*, 6, 177-208.

Klein, R.G., Cruz-Uribe, K. (1987). Large mammal and tortoise bones from Eland's Bay Cave and nearby sites, Western Cape Province, South Africa. In: Parkington, J., Hall, M. (Eds). *Papers in the Prehistory of the Western Cape*. *BAR International Series*, 332, 132-163.

Knight, H. (1970) Assemblage from Ringaringa, Stewart Isalnd. *New Zealand Archaeological Association Newsletter*, 76-83.

Li H., Durbin, R. (2009) Fast and accurate short read alignment with Burrows-Wheeler Transform. *Bioinformatics*, 25, 1754–1760.

Li, H., Handsaker, B., Wysoker, A., Fennell, T., Ruan, J., Homer, N., Marth, G., Abecasis, G., Durbin, R., 1000 Genome Project Data Processing Subgroup. (2009) The Sequence alignment/map (SAM) format and SAMtools, *Bioinformatics*, 25, 2078–2079.

Ling, J.K. (1999a) Elephant seal oil cargoes from King Island, Bass Strait, 1802-1819: with estimates of numbers killed and size of the original population. *Papers and Proceedings of the Royal Society of Tasmania,* pp 51-56.

Ling, J.K., Bryden, M.M. (1982) *Mirounga leonina*. *Mammalian Species*, 391, 1-8.

Magoč, T., Salzberg, S.L. (2011) FLASH: fast length adjustment of short reads to improve genome assemblies. *Bioinformatics*, 27, 2957-2963.

Maricic, T., Whitten, M., Pääbo, S. (2010) Multiplexed DNA sequence capture of mitochondrial genomes using PCR products. *PLoS ONE*, 5, e14004.
Martin, M. (2011) Cutadapt removes adapter sequences from high-throughput sequencing reads. *EMBnet Journal*, 17, 10-12.
Meyer,M., Kircher M. (2010) Illumina sequencing library preparation for highly multiplexed target capture and sequencing. *Cold Spring Harbour Protocols. 2010* (6).
Nagaoka, L. (2000). *Resource Depression, Extinction, and Subsistence Change in Prehistoric Southern New Zealand*. University of Washington.
Peck, D. R., Congdon, B.C. 2004. Reconciling historical processes and population structure in the sooty tern *Sterna fuscata*. *Journal of Avian Biology,* 35**,** 327–335.
Petchey, F., Anderson, A., Hogg, A.G. (2008). The marine reservoir effect in the Southern Ocean : an evaluation of extant and new R values and their application to archaeological chronologies. *Journal of the Royal Society of New Zealand*, 38, 243–262.
Prost, S., Anderson, C.N.K. (2011) TempNet: a method to display statistical parsimony networks for heterochronous DNA sequence data. *Methods in Ecology and Evolution,* 2, 663–667.
Rohland, N., Siedel, H., Hofreiter, M. (2010) A rapid column-based ancient DNA extraction method for increased sample throughput. *Molecular Ecology Resources*, 10, 677–683.
Rozas, J., Ferrer-Mata, A., Sánchez-DelBarrio, J. C., Guirao-Rico, S., Librado, P., Ramos-Onsins, S. E., Sánchez-Gracia, A. (2017) DnaSP 6: DNA sequence polymorphism analysis of large data sets. *Molecular Biology and Evolution,* 34**,** 3299–3302.
Schweitzer, F.R. (1979). Excavations at Die Kelders, Cape Province, South Africa: the Holocene deposits. *Annals of the South African Museum*, 78, 101-233.
Seersholm, F.V., Cole, T.L., Grealy, A., Bunce, M. (2018) Subsistence practices, past biodiversity, and anthropogenic impacts revealed by New Zealand-wide ancient DNA survey. *Proceedings of the National Academy of Sciences USA*, 115, 7771-7776.
Slade, R. W., Moritz, C., Hoelzel, A.R., Burton, H.R. (1998) Molecular population genetics of the southern elephant seal *Mirounga leonina*. *Genetics,* 149**,** 1945–1957.
Smith, I. (1985) *Sea mammal hunting and prehistoric subsistence in New Zealand.* University of Otago. PhD Thesis.
Smith, I. (1999) Settlement permanence and function at Pleasant River mouth, East Otago, New Zealand. *New Zealand Journal of Archaeology*, 19, 27-79.
Smith, I. (2013). Pre-European Maori exploitation of marine resources in two New Zealand case study areas: species range and temporal change. *Journal of the Royal Society of New Zealand*, 43, 1–37.
Smith, I., Wernham, P. (1976) Survey of archaeological sites: Te Awapatiki to Hapupu, Hanson Bay, Chatham Island. Working Papers in Chatham Islands Archaeology, 1, Anthropology Department, University of Otago.
Stockton, J. (1982) Seals in Tasmanian prehistory. *Proceedings of the Royal Society of Victoria*, 94, 53-60.
Sutton, D.G., Marshall, Y.M. (1980) Coastal hunting in the subantarctic zone. *New Zealand Journal of Archaeology*, 2, 25-49.

Valenzuela-Toro, A.M., Gutstein, C.S., Suárez, M.E., Otero, R., Pyenson, N.D., (2015). Elephant seal (*Mirounga* sp.) from the Pleistocene of the Antofagasta Region, northern Chile. *Journal of Vertebrate Paleontology*, 35, e918883.

Verry, A.J.F., Mitchell, K.J., Rawlence, N.J. (2022). Genetic evidence for post-glacial expansion from a southern refugium in the eastern moa (*Emeus crassus*). *Biology Letters*, 18, 20220013.

Walsh, P., Metzger, D., Higuchi, R. (1991) Chelex 100 as a medium for simple extraction of DNA for PCR-based typing from forensic material. *BioTechniques*, 10, 506–513.

Warneke, R.M. (1976) Preliminary report on the distribution and abundance of seals in the Australian region. *Scientific Consultation on Marine Mammals, Food and Agriculture Organisation of the United Nations, Bergen, Norway.*
